# Stromal Oncostatin M axis promotes breast cancer progression

**DOI:** 10.1101/2020.10.30.356774

**Authors:** Angela M. Araujo, Andrea Abaurrea, Peio Azcoaga, Joanna I. López-Velazco, Ricardo Rezola, Iñaki Osorio-Querejeta, Fátima Valdés-Mora, Juana M. Flores, Liam Jenkins, Patricia Fernández-Nogueira, Nicola Ferrari, Natalia Martín-Martín, Alexandar Tzankov, Serenella Eppenberger-Castori, Isabel Alvarez-Lopez, Ander Urruticoechea, Paloma Bragado, Nicholas Coleman, Arkaitz Carracedo, David Gallego-Ortega, Fernando Calvo, Clare M. Isacke, Maria M. Caffarel, Charles H. Lawrie

## Abstract

Cancer cells are constantly communicating with the surrounding tumour microenvironment (TME) and they hijack physiological cell interactions to overcome immune system surveillance and promote cancer progression^1,2^. However, the contribution of stromal cells to the reprogramming of the TME is not well understood. In this study we provide unprecedented evidence of the role of the cytokine Oncostatin M (OSM) as central node for multicellular interactions between immune and non-immune stroma and the epithelial compartment. We show that stromal expression of the OSM:Oncostatin M Receptor (OSMR) axis plays a key role in breast cancer progression. OSMR deletion in a multistage breast cancer model delays tumour onset, tumour growth and reduces metastatic burden. We ascribed causality to the stromal function of OSM axis by demonstrating reduced tumour burden of syngeneic tumours implanted in mice. Single-cell and bioinformatic analysis of murine and human breast tumours revealed that the expression of OSM signalling components is compartmentalized in the tumour stroma. OSM expression is restricted to myeloid cells, whereas OSMR expression is detected predominantly in fibroblasts and, to a lower extent, cancer cells. Myeloid-derived OSM reprograms fibroblasts to a more contractile and pro-tumorigenic phenotype, elicits the secretion of VEGF and pro-inflammatory chemokines (e.g. CXCL1 and CXCL16), leading to increased neutrophil and macrophage recruitment. In summary, our work sheds light on the mechanism of immune regulation by the tumour microenvironment, and supports that targeting OSM:OSMR interactions is a potential therapeutic strategy to inhibit tumour-promoting inflammation and breast cancer progression.

## Main text

The tumour microenvironment (TME), composed by different cell types (e.g. fibroblasts, adipocytes, endothelial and infiltrating immune cells), harbours complex cell interactions that are often manipulated and hijacked by tumour cells in with every step of cancer progression^2^. However, the contribution of stromal cells to the reprogramming of the tumour microenvironment is poorly understood. Here, we discovered that the cytokine Oncostatin M (OSM) acts as a central regulator of the crosstalk between immune stroma and non-immune stroma, favouring breast cancer progression and metastasis. First, we set to study the contribution of OSM signalling in the genetic mouse model MMTV-PyMT, widely used to study breast cancer progression in a fully competent tumour microenvironment and immune system^3^. We crossed Oncostatin M Receptor (OSMR) deficient mice with MMTV-PyMT as illustrated by the experiment scheme in Fig. 1a. MMTV-PyMT: OSMR KO females showed a significant delay in tumour onset, tumour growth and a reduced tumour burden at 14 weeks of age (Fig. 1b-d and Extended Data Fig. 1a-c). Importantly, OSMR deletion also reduced the malignancy of the tumours, assessed by histopathological analysis, as it reduced the percentage of mice with malignant carcinomas and increased the proportion of mice with pre-malignant adenomas/ mammary intraepithelial neoplasia (MIN) or no tumours (Fig. 1e and Extended Data Fig. 1d, P value = 0,007 for Chi Square test comparing malignant lesions versus pre-malignant lesions or no lesions). Interestingly, when compared to their controls, tumours in OSMR-deficient mice showed decreased levels of the extracellular matrix protein fibronectin, predominantly produced by CAFs^4^ (Fig 1f), increased levels of apoptosis, but similar degree of proliferation (Extended Data Fig. 1e). Finally, OSMR deficiency produced a remarkable reduction in the percentage of animals with lung metastases (Fig. 1g,h).

**Fig. 1:**
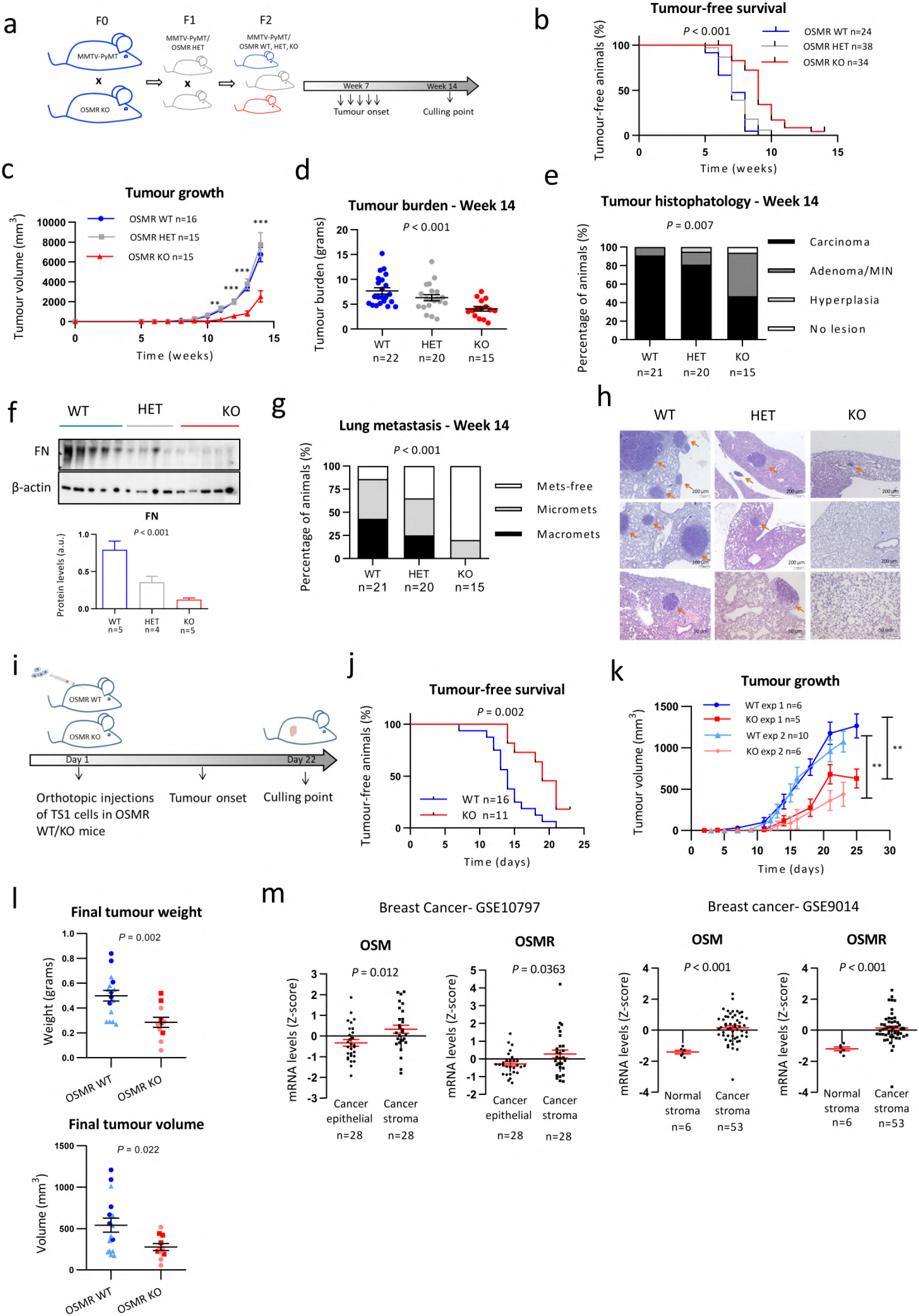
Stromal OSM:OSMR axis promotes breast cancer progression. **a)** Experimental set-up of the *in vivo* experiment designed to assess the importance of OSMR signalling in disease progression of the MMTV-PyMT mouse model. **b-d)** Kaplan-Meier curves for tumour-free survival **(b),** tumour growth **(c)** and final tumour burden **(d)** in MMTV-PyMT:OSMR wild −t ype (WT), M MTV-PyM T:OSM R heterozygous (HET), and MMTV-PyMT:OSMR knockout (KO) mice. **e)** Histopathological analysis of tumours at week 14. Graph represents percentage of mice bearing carcinomas, adenomas, hyperplasia and no lesions in mammary glands. P value was determined comparing number of mice with malignant carcinoma *vs.* non-malignant phenotypes (Adenoma, hyperplasia) and no lesions using Chi-square test. **f)** West ern blot (upper panel) and densitometric analysis (lower panel) of fibronectin protein levels in tumours at week 14 from animals of the different genotypes. **g)** Percentage of animals with lung metastases at 14 weeks of age. Lung tumour masses were classified as macrometastases when they were visible at dissection and as micrometastases when they were only detectable by hematoxylin and eosin staining. P value was determined comparing animals with metastasis (macro and micro) vs. non-metastasis using Chi-square test. **h)** Representative pictures of lung metastases at week 14 in OSMR WT, HET and KO animals. **i)** Experimental set-up of the *in vivo* experiment designed to assess the importance of OSMR signalling in the tumour microenvironment, in which TSl cells were orthotopically injected into the mammary fat pad of OSMR WT and KO mice. **j-1)** Kaplan-Meier curves for tumour-free survival **(j),** tumour growth **(k)** and final tumour volume and weight after dissection **(I)** of orthotopic tumours described in i). Two independent experiments were performed, and the results were combined in **j** and **I. m)** OSM and OSMR mRNA expression in paired cancer epithelial vs. cancer stroma (left) and normal stroma vs. cancer stroma breast cancer samples (right). Data were downloaded from GEO Datasets (GSE10797 and GSE9014). In **b,j)** P values were calculated using the M ant el-Co x test; in **c,k,l,m)** P values were determined using unpaired two-tailed t test; ind, **f)** P values were determined using one-way ANOVA t est. Unless otherwise specified graphs represent mean ± SEM ** p < 0,01; *** p < 0,001.

These results show that OSM signalling is causally associated with tumour aggressiveness but, surprisingly, by using syngeneic cancer models, we found that this association requires, at least in part, the presence of the OSM:OSMR axis in the tumour stroma. We injected TS1 cells, derived from a MMTV-PyMT tumour^5^, orthotopically into the mammary gland of syngeneic OSMR deficient (KO) and wild-type (WT) control mice (Fig. 1i). This model allows the assessment of the contribution of stromal OSMR signalling to cancer progression as OSMR is only depleted in the tumour microenvironment while TS1 cancer cells express OSMR that can be activated by host-derived OSM (Extended Data Fig. 1f,g). Depletion of OSMR in the tumour microenvironment resulted in delayed tumour onset and tumour growth (Fig. 1j-l and Extended Data Fig. 1g) confirming that stromal OSMR signalling contributes to cancer progression.

Analysis of published gene expression profiles of breast cancer demonstrated that both OSM and OSMR are increased in human breast cancer stroma, compared to cancer epithelial compartment and healthy stroma (Fig. 1m). A similar pattern of OSM:OSMR expression was observed in other cancer types including colorectal and ovarian cancers (Extended Data Fig. 1h). We also observed that increased OSM mRNA levels associated with decreased disease-free survival (Extended Data Fig.1i) in the Metabric^6^ and Wang^7^ breast cancer datasets. Analysis of TCGA data by Kaplan-Meier Plotter^8^ showed that high OSM levels were significantly associated with worse overall survival in other cancer types (Extended Data Fig. 1j).

As we found an unexpected contribution of stromal OSM: OSMR axis to breast cancer progression, we performed single cell RNA-seq analysis of mammary tumours from the MMTV-PyMT model to decipher which cells were responsible to produce OSM and to express OSMR in the breast cancer context (Fig. 2a). Our data indicate that the ligand OSM is almost exclusively expressed by the myeloid cell population, while the receptor OSMR is mainly expressed by the fibroblasts and some of the cancer epithelial clusters (Fig. 2a-c). The OSM:OSMR signaling module exhibits a distinct microenvironment-restricted expression and it differs from the one observed for other cytokine-receptor pairs of the same family such as IL6:IL6R (Il6ra) and LIF:LIFR (Fig. 2b,c), supporting that OSM exerts distinct and unique functions from other members of the family^9^. Il6st (GP130) is the common subunit receptor for OSM, IL6, LIF and other cytokines of the family and is ubiquitously expressed (Fig. 2b,c), being the expression of the other receptor subunits more restricted and tightly regulated. RT-qPCR analysis of FACS-sorted breast TS1 orthotopic tumours^5^ confirmed expression of OSM in the myeloid population and expression of OSMR in fibroblasts and in cancer cells (Extended Data Fig. 2a and Supplementary Fig. 1). Similar results were obtained when analysing FACS sorted populations of MMTV-PyMT tumours (Extended Data Fig. 2b). An identical pattern of OSM:OSMR expression is maintained in the human setting, as demonstrated by RT-qPCR quantification in a large panel of human cell lines and analysis of RNA-seq data from the Human Protein Atlas^10^ (Fig. 2d and Extended Data Fig. 2c,d). OSM mRNA expression was restricted to undifferentiated and macrophage-like differentiated HL-60 cells^11^ (Extended Data Fig.2c) and lymphoid and myeloid cell lines (Extended Data Fig. 2d). Conversely, OSMR was only detected by RT-qPCR in breast cancer cells and fibroblasts, showing significantly higher expression in fibroblasts compared to epithelial cells (Fig. 2d and Extended Data Fig. 2c). Analysis of a battery of human cell lines^10^ confirmed expression of OSMR only in epithelial, endothelial and fibroblast cell lines and not in immune cell lines (Extended Data Fig. 2d). To prove the relevance of our previous findings in human cancer clinical data, we used the TIMER^12^ and xCell^13^ web resources to analyse the association between OSM and OSMR expression and TME composition in two different clinical breast cancer datasets^8,14^. TIMER analysis showed that OSMR mRNA expression significantly correlates with fibroblast enrichment in human breast cancer, while OSM mRNA levels show the most significant associations with myeloid macrophage and neutrophil signatures (Fig. 2e). This analysis also showed that OSM and OSMR mRNA expression inversely correlated with tumour cell purity. The OSMR and OSM associations with fibroblasts and myeloid cell infiltration respectively, were validated by xCell in a different clinical dataset (Fig. 2f). A similar pattern of OSM:OSMR expression was observed in FACS-sorted colorectal tumours (Extended data Fig. 2e). Altogether, our data reveal that OSM and OSMR are stroma-expressed molecules, and point to paracrine OSM:OSMR signalling in cancer, as ligand and receptor are expressed by different cell types in the tumour microenvironment.

**Fig. 2:**
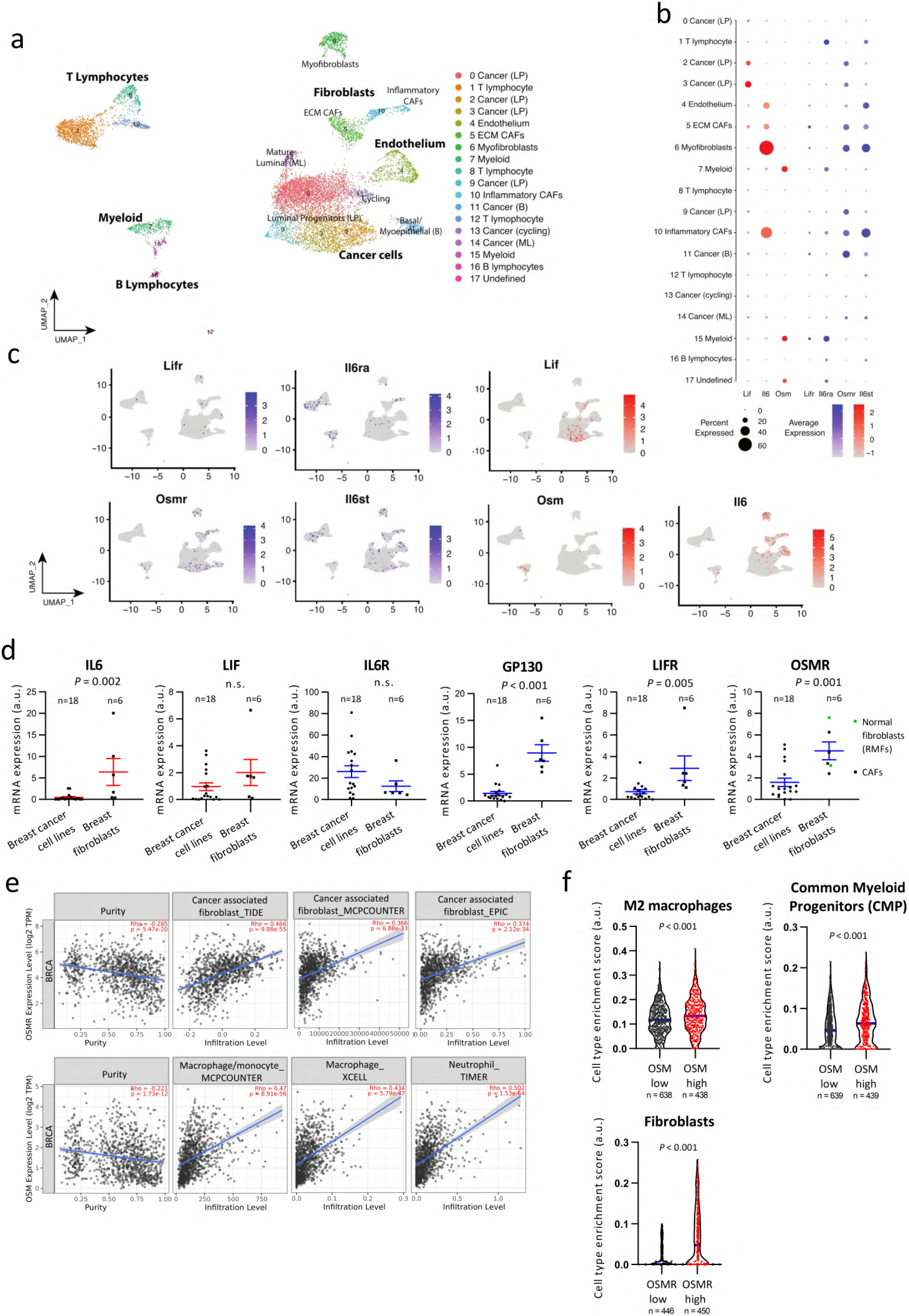
The OSM:OSMR signalling module exhibits a distinct microenvironment-restricted expression. **a)** UMAP plot showing cell clusters defined in each of the main cell lineages. Far right column depicts the main cell lineage of origin for each cluster, showing 7 clusters of epithelial origin, 6 immune and 4 stromal. LP: luminal progenitors, ECM: extracellular matrix, B: basal, ML: mature luminal. **b)** Dot plot representing the expression level (red or blue jet) and the number of expressing cells (dot size) of the indicated genes in each cluster. **c)** Feature UMAP plots showing the expression of the indicated genes in each of the main cell clusters. **d)** mRNA expression levels of the indicated IL-6 family members and associated receptors analysed by RT-qPCR in a panel of breast cancer cell lines and immortalized fibroblasts. In the OSMR graph (right panel), green and dark dots represent normal mammary fibroblasts and CAFs, respectively. Graphs represent mean± SEM . P values were determined using the unpaired two-tailed t test. n.s. non-significant. RMF: reduction mammoplasty fibroblasts. **e)** Correlation of OSMR (upper panels) and OSM (lower panels) expression with tumour purity and infiltration level of indicated cell types in breast cancer samples. Data were downloaded from TIMER web platform. Spearman correlation coefficients and P values are shown. **f)** Truncated violin plots showing cell type enrichment of the indicated populations in breast tumours according to high (top quartile) or low (lower quartile) OSM or OSMR expression. Data were obtained using xCell web resource on 1809 breast cancer samples from Kaplan-Meier Plotter website. P values were determined using Mann-Whitney’s test.

As we previously observed that fibroblasts were the cell population with higher levels of OSMR within the tumour (Fig. 2 and Extended data Fig. 2), we performed complementary *in vitro* and *in vivo* experiments to assess the effect of OSMR activation in mammary cancer-associated and normal fibroblasts derived from human breast tumours and reduction mammoplasty surgeries respectively^15^. The ability to remodel the extracellular matrix is a hallmark of CAFs^4^. Importantly, OSM treatment enhanced the capacity of CAFs (CAF-173 and CAF-318) to contract collagen matrices and, interestingly, the effect was not observed in non-cancerous skin and breast fibroblasts (HS27 and RMF-31, respectively) (Fig. 3a). To further investigate the role of OSM in potentiating CAFs activation, we selected RMF-31 to be used as a model of normal breast fibroblasts and CAF-173 as a model of CAFs. In accordance with the contractility experiments, OSM promoted the growth of 3D CAF spheroids while it did not affect normal mammary fibroblasts 3D spheroids (Fig. 3b). Similarly, OSM induced the expression of classical CAF markers such as FAP, POSTN, VEGF and IL6^4^, only in CAF-173 CAFs, and not in normal RMF-31 fibroblasts (Fig. 3c). Of interest, OSMR was similarly expressed in normal and cancer fibroblasts (Fig. 2d) and the pathway was functional in both CAFs and normal fibroblasts, as suggested by OSM induction of OSMR expression in both cell lines (Extended Data Fig. 3a), classical hallmark of OSMR activation^16^. Gene set enrichment analysis (GSEA) of transcriptomic data of CAF-173 treated with OSM or vehicle, showed that OSM induced signatures related to fibroblast activation and JAK-STAT3 signalling, in agreement with increased STAT3 phosphorylation by OSM (Fig. 3d and Extended Data Fig. 3b,c). A transcriptional signature composed by the top differentially expressed genes by OSM in CAF-173 was enriched in the breast cancer stroma GSE9014 dataset compared to normal stroma (Extended Data Fig. 3d). Importantly, the top 4 genes induced by OSM in CAF-173 (SERPINB4, THBS1, RARRES1 and TNC; Extended Data Fig. 3e) are associated with decreased recurrence-free survival in breast cancer patients (Fig. 3e). In addition, THBS1, RARRES1 and TNC levels correlate with OSMR expression in breast cancer clinical samples (Extended Data Fig. 3f). These results indicate that OSM induces in CAFs the expression of pro-malignant genes, including fibroblast activation markers and genes associated with JAK-STAT3 signalling.

**Fig. 3:**
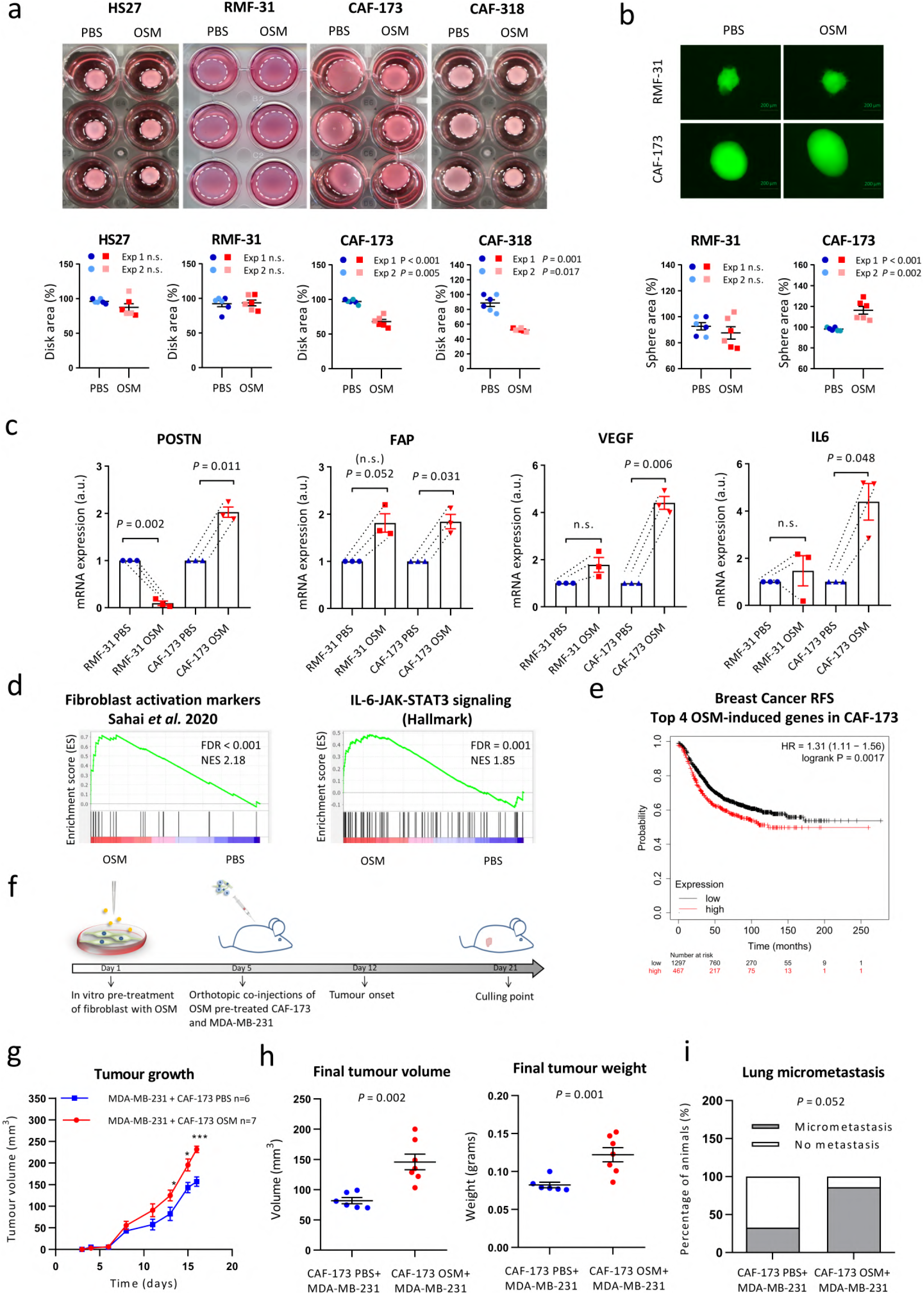
OSM activates cancer-associated fibroblasts (CAFs) *in vitro* and *in vivo* promoting tumour progression. **a)** Representative pictures of collagen contraction assays (upper panels) and quantification of collagen disk areas (lower panels) of fibroblasts pre-treated in monolayer with PBS or OSM and seeded in collagen. **b)** Representative pictures (upper panels) and area quantification (lower panels) of 3D spheres proliferation assays of fibroblasts treated with PBS or OSM. In **a** and **b,** graphs represent mean± SEM, and 2 independent experiments are plotted. **c)** RT-qPCR analysis of mRNA levels of activation markers in normal (RMF-31) and cancer-associated (CAF-173) fibroblasts cultured in 3D with PBS or OSM. Graphs represent mean± SEM of 3 independent experiments. P values were determined using paired two-tailed t tests. **d)** GSEA showing enrichment of the indicated signatures in microarray data of CAF-173 treated with OSM. **e)** Kaplan-Meier curves showing relapse-free survival (RFS) for breast cancer samples according to the high or low expression of top 4 genes induced by OSM in CAF-173. Data were obtained using **KM** plotter website. **f)** Experimental set up of the *in vivo* experiment designed to assess the contribution of OSMR activation in fibroblasts to cancer progression. CAF-173 were pre-treated with OSM or PBS for 4 days prior to injection and were co­ injected with MDA-MB-231 (500.000 cells each cell line) in matrigel (1:1ratio) in the mammary gland fat pad of nude mice. n=6 animals with MDA-MB-231+ CAF-173 PBS cells injected; and n=7 animals with MDA-M B-231+ CAF-173 OSM cells injected. **g,h)** Tumour growth **(g)** and final tumour volume and weight after dissection **(h)** of orthotopic tumours described in **f).** Graphs represent mean± SEM. **i)** Percentage of animals with lung micrometastasis assessed using qPCR analysis of genomic human ALU sequences. Graph represents the percentage of animals with detectable qPCR signal and P value was calculated using the Chi-square test. n.s. non-significant.* p < 0,05; *** p < 0,001. P values were calculated using the unpaired two-tailed t test unless specified.

In order to test if the OSM-induced changes in CAFs contributed to breast cancer progression *in vivo*, we pre-treated CAF-173 CAFs with OSM or vehicle for 4 days *in vitro* and orthotopically co-injected them with MDA-MB-231 breast cancer cells into Athymic Nude-Foxn1nu mice as described in the experiment timeline in Fig. 3f. Activation of fibroblasts by OSM promoted tumour growth (Fig. 3g,h) and exhibited a trend to increase lung colonization (Fig. 3i), assessed by qPCR analysis of human Alu DNA sequences in the lungs^17^. Conversely, OSMR downregulation by shRNA in CAF-173 delayed tumour onset and tumour growth at early stages when co-injected with MDA-MB-231 breast cancer cells ectopically expressing human OSM (Extended Data Fig. 3g-j). In addition, downregulation of OSMR in CAFs decreased IL6 expression in tumours, suggesting that OSM is inducing the expression of similar targets *in vivo* (Extended Data Fig.3k). Moreover, the tumours with OSMR silencing in CAFs showed reduced levels of GFP (Extended Data Fig. 3k), suggesting reduced levels of CAFs in this experimental group, probably due to impaired CAF proliferation upon OSMR reduction, in line with the increased size of CAF spheres observed after OSMR activation (Fig. 3b). Together, our data prove that OSM:OSMR signalling activates CAFs and that this contributes to cancer progression.

In an attempt to understand how OSMR activation in the stroma was inducing malignancy we deepened into our transcriptomic data of CAFs (CAF-173) treated with OSM. Microarray data indicated that pathways and signatures related to leukocyte chemotaxis and inflammatory response were significantly enriched by OSM (Fig. 4a,b). Interestingly, transcriptomic analysis of breast cancer cells (MDA-MB-231) activated by OSM showed enrichment of similar pathways (Extended Data Fig. 4a,b). These data suggested that, upon OSMR activation by OSM, both CAFs and cancer cells could be involved in shaping the tumour microenvironment by recruiting leukocytes to the tumour site. Analysis of a panel of 31 chemokines by antibody array showed that OSM induced expression of important chemoattractants (Fig. 4c and Extended Data Fig. 4c,d). Some of these factors were exclusive of CAFs (mainly CXCL10 and CXCL12), others of cancer cells (mainly CXCL7 and CCL20) and some factors, such as CCL2, were common for both cell types. Vascular endothelial growth factor (VEGF) can also modulate tumour immunity by inducing macrophage and myeloid-derived suppressor cells (MDSCs) recruitment^18^ and we previously showed that it is an OSMR target^16^. As seen in Fig. 4d and Extended Data Fig. 4e, VEGF levels were increased upon OSM treatment both in CAFs and tumour cells. As some of the OSM-induced chemokines are potent myeloid chemoattractants (e.g. VEGF, CCL2, CXCL12)^19,20^, we sought to determine whether myeloid cell populations were altered in tumours after OSMR signalling abrogation. We performed immunostaining of macrophages and neutrophils, assessed by F4/80 and Ly6G positivity respectively^2,21,22^, and we observed that these two populations were reduced in MMTV-PyMT: OSMR KO tumours compared to OSMR WT tumours (Fig. 4e). Interestingly, VEGF, CXCL1 and CXCL16 levels were reduced in the serum of tumour bearing MMTV-PyMT: OSMR KO mice compared to control mice (Fig. 4f), all factors being involved in myeloid cell recruitment^18,23,24^. In summary, these results show that OSM:OSMR signalling in stroma and cancer cells induces cytokine secretion and myeloid cell recruitment. Our findings are clinically relevant as VEGF, CXCL1 and CXCL16 mRNA expression is associated with decreased recurrence-free survival and with OSM and/or OSMR levels in breast cancer patients (Fig. 4g,h). As OSM is mainly expressed by myeloid cells (Fig. 2 and Extended Data Fig. 2), our data point to the existence of a feedback positive loop where OSM signalling in both CAFs and cancer cells induces the recruitment of more myeloid cells which will in turn secrete more OSM within the tumour. Moreover, conditioned media from cancer cells pre-treated with OSM further increased OSM expression in macrophage-like differentiated HL-60 cells (Extended Data Fig. 4f). We did not observe this effect with conditioned media from OSM-activated CAFs or with OSM itself. These results suggest that OSMR activation in cancer cells, not only increases the secretion of chemokines involved in myeloid cell recruitment, but also induces OSM expression in these myeloid cells, doubly contributing to the aforementioned feed-forward loop.

**Fig. 4:**
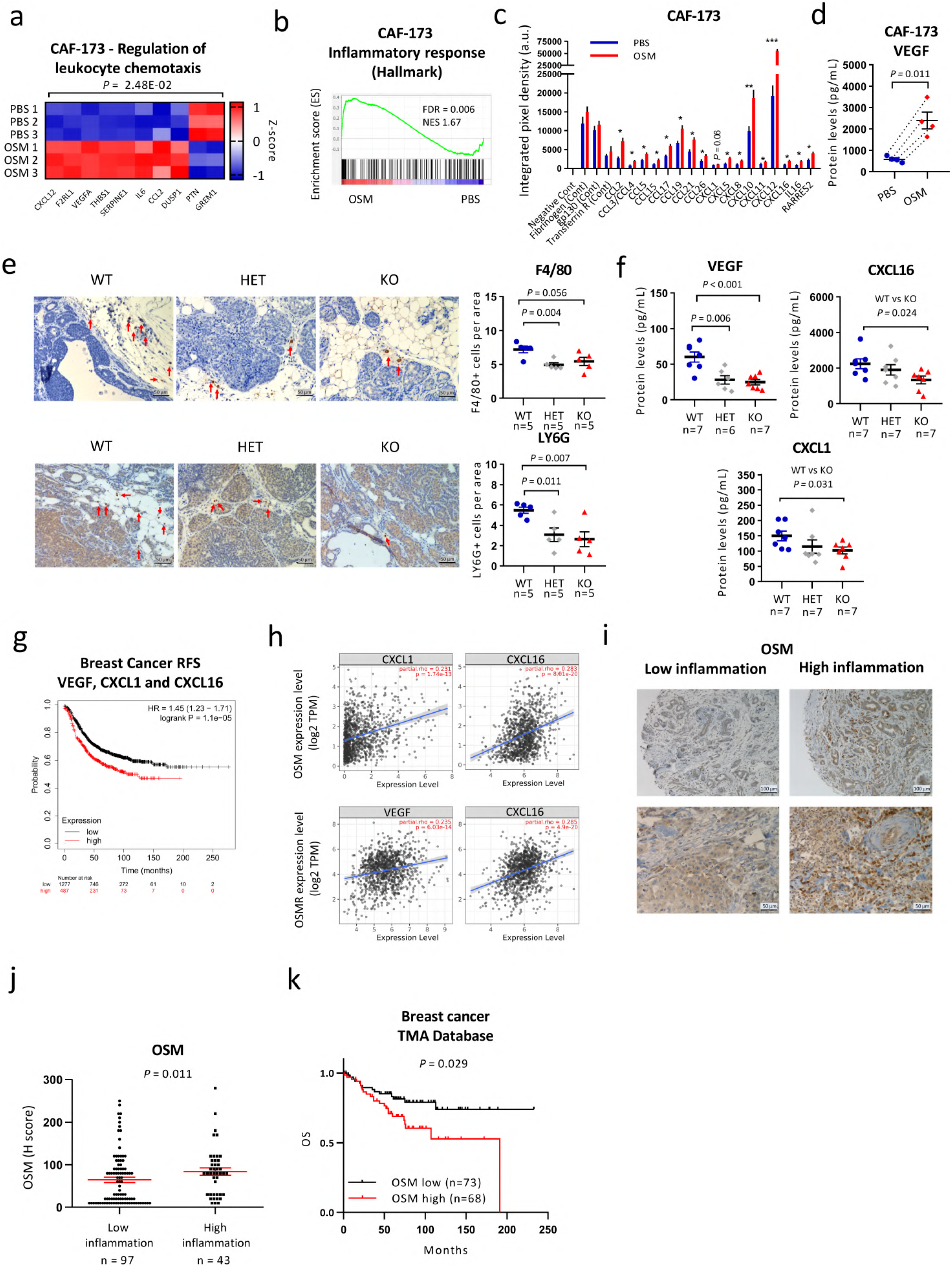
OSM:OSMR signalling in CAFs induces cytokine secretion and myeloid recruitment. **a)** Heatmap showing normalized mRNA expression of genes induced by OSM in CAF-173 and included in the indicated GO pathway. **b)** GSEA showing enrichment of inflammatory hallmark signature in microarray expression data of CAF-173 spheres treated with OSM 30ng/ml for 4 days. **c,d)** Chemokine array analysis **(c)** and VEGF levels **(d)** of conditioned media from CAF-173 treated with PBS (Control) or OSM 30ng/ml for 72 hours. Graphs represent mean± SEM (n=4 independent experiments). P values were determined using paired two-tailed t tests. **e)** Representative pictures (left panels) and quantification (right panels) of F4/80 (upper panels) and LY6G (lower panels) immunohistochemistry staining in tumours from MMTV-PyM T:OSM R wild −typ e (WT), heterozygous (HET), and KO mice at 14 weeks of age. Quantification was performed by manual counting of positive cells per area in a total of 8 pictures per tumour and 5 tumours per group. **f)** VEGF, CXCLl and CXCL16 levels in plasma from MMTV-PyMT: OSMR WT, HET and KO mice at 14 weeks of age (n=S per group) analysed by Luminex assay. In **e,f)** graphs represent mean± SEM . P values between the different groups were determined using unpaired two tailed t tests. **g)** Kaplan-Meier curves showing relapse-free survival (RFS) for breast cancer samples according to the expre ssion of VEGF, CXCLl and CXCL16. Data were downloaded from KM plotter. **h)** Correlation of OSM (top) and OSMR levels (bottom panel) with VEGF, CXCLl and CXCL16 expression in breast cancer samples. Data were downloaded from TIMER web platform. Spearman correlation coefficients and P values are shown. **i, j)** Representative pictures **(i)** and quantificat ion **(j)** of OSM staining in samples for breast cancer patients with high and low inflammation. Graph represents mean± SEM and P value was determined using Mann-Whitney’s test. **k)** Kaplan-M eier curves showing overall survival (OS) for breast cancer patients analyised in **i** and **j,** with high vs. low OSM expression. In **g** and **k,** P values were determined using the Mantel-Cox test and high and low expression levels were determined by automatic cutoff.

Analysis of OSM protein levels in 141 samples of early breast cancer samples confirmed the association between OSM expression and increased inflammation in a clinical setting (Fig. 4i,j). Inflammation was assessed by the pathologist as infiltration of inflammatory cells from all lymphoid and myeloid subtypes. We observed that OSM was mainly expressed by myeloid-like cells as determined by their larger size and more irregular shape (Fig. 4i). Lymphoid cells, characterized by being smaller and round and by having a round nucleus with little cytoplasm, showed very low or negative OSM expression (Fig. 4i). Importantly, high OSM protein levels were associated with decreased overall survival in this dataset (P=0.029, Fig. 4k).

Cytokines are important players in inflammation, a process associated with tumour progression^1^. Even the cancers not directly associated with persistent infections or chronic inflammation, such as breast cancer, exhibit tumour-elicited inflammation, which has important consequences in tumour promotion, progression and metastasis^25,26^. Understanding how inflammatory signals orchestrate pro-malignant effects in the different cell compartments within the tumour microenvironment is key to design new therapeutic strategies to target tumour-promoting inflammation. Our study identifies the pro-inflammatory cytokine OSM as a crucial mediator of the crosstalk between different cell types within the tumour by activating an intriguing pro-tumoural “*ménage-à-trois*” between myeloid cells, CAFs and cancer cells (Extended Data Fig. 5). OSM:OSMR interactions could be blocked by antibody based inhibition, a strategy that has had a major impact on cancer^27^, which makes them a promising candidate for therapeutic targeting. Interestingly, anti-OSM humanized antibodies have proven to be safe and well tolerated^28^ and are now in Phase 2 clinical trials for the treatment of inflammatory diseases, such as systemic sclerosis and Crohn’s disease. Together, our findings further strengthen the case for the pre-clinical investigation of OSM:OSMR blocking antibodies as a targeted anti-cancer therapy.

## Methods

### Tissue Microarrays

Formalin-fixed and paraffin-embedded blocks of 141 tumour tissues from cases surgically resected at the University Hospital Basel between 1991 and 2013, and included in tissue microarrays (TMAs), were used for analysis of OSM protein expression in human samples. All patients have given informed consent for their archival tissue to be used for scientific research, and the TMA construction was authorized by the Hospital Ethic Committee. TMAs were generated by punching 1 mm spot of each sample. Complete histopathological information, date and cause of death, as well as date of local and/or distant relapse were available for all the patients. Tissue sections were subjected to a heat-induced antigen retrieval step prior to exposure to primary OSM antibody (1:50, HPA029814, Sigma-Aldrich). Immunodetection was performed using the Roche Ventana BenchMark ULTRA IHC/ISH staining system, with DAB as the chromogen. Cases were reviewed by two independent pathologists and OSM staining was evaluated by the semiquantitative method of H-score (or “histo” score), used to assess the extent of immunoreactivity in tumour samples^29^. Inflammation was assessed by a pathologist as high or low tumour infiltration of immune cells according to their morphology.

### Gene expression analysis of clinical datasets and bioinformatics analysis

Disease-free survival (DFS) of patients based on OSM mRNA expression was calculated using data from the publicly available METABRIC^6^ and Wang^7^ datasets with the CANCERTOOL interface^30^. Kaplan-Meier curves showing overall (OS) or relapse-free survival (RFS) of patients from various cancer types according to the expression of different genes by RNA-seq-analysis were obtained from Kaplan-Meier Plotter website^8^. Best threshold cutoffs were selected automatically by the program. RNA-seq data from 64 cell lines was retrieved from the Human ProteinAtlas^10^. RNA consensus normalized expression values (NX) were plotted for OSM and OSMR transcripts using GraphPad software. Associations between OSMR and OSM mRNA expression and enrichment of different cell types were analysed by using xCell^13^ on 1809 breast cancer samples from Kaplan-Meier Plotter website^8^ and TIMER2.0 which incorporates 1100 breast cancer samples from TCGA^12^. TIMER2.0 was also used to analyse gene expression correlations, after purity adjustment. All correlations were calculated with the Spearman’s rank correlation coefficient. Gene expression analyses of human tumour stroma and epithelia were retrieved from NCBI Gene Expression Omnibus (GEO): Finak (GSE9014, Breast)^31^, Casey (GSE10797, Breast)^32^, Yeung (GSE40595, Ovary)^33^, Nishida (GSE35602, Colon)^34^, and Calon (GSE39396, Colon)^35^. For Affimetrix-based arrays, probe-to-gene mapping was performed using Jetset, while for the rest, highest variance probes were selected. Unless otherwise stated, expression values for each gene were z-score normalized. GO analysis was performed using Panther^36^.

### Single-cell RNA sequencing (scRNA-seq)

Drop-seq dataset^37^ raw data for MMTV-PyMT (WT) tumours were obtained from Valdes-Mora *et al. (2020)*^38^. This subset was subsequently analysed using Seurat^39^ (v Seurat 3.2). Briefly, a total of 9,636 sequenced cells from 8 MMTV-PyMT tumours pass the QC filter, with <5% mitochondrial to nuclear gene content^37^, and <8,000 molecules/cell as they potentially represented cell doublets. Downstream analysis was performed according to Butler *et al.* (2018)^39^, using 30 principal components to build a Shared Nearest Neighbour (SNN) graph calculating k-nearest neighbour (Jaccard Index) for each cell, subsequent cluster calling and UMAP dimensional reduction projection^40^.

### Cell culture

Human breast cancer-associated (CAF-173, CAF-200, CAF-220 and CAF-318) and normal (RMF-31 and RMF-39) fibroblasts were derived from human breast tumours and reduction mammoplasty surgeries respectively, immortalized, tagged with GFP and cultured in collagen pre-coated flasks^15^. The aforementioned human mammary fibroblasts, TS1 cells derived from primary tumours of the MMTV-PYMT mice model^5,41^, LM2 breast cancer cells (kindly donated by Dr Roger Gomis) and HS27 skin fibroblasts (kindly donated by Dr Ander Izeta) were cultured in DMEM medium supplemented with 10% fetal bovine serum (FBS), 1% glutamine, and 1% penicillin and streptomycin. HL-60 promyeloblast cell line, the human embryonic kidney cell line HEK293T and human breast cancer cell lines (MDA-MB-231, BT-549, HCC38, MDA-MB-157, SUM149PT, HCC1806, HCC70, MDA-MB-468, HCC1569, HCC1954, SK-BR-3, MDA-MB-453, CAMA-1, ZR-75-1, T47D, MCF-7, BT-474) were purchased from American Type Culture Collection (ATCC) and cultured following ATCC instructions. All cell lines were authenticated by short tandem repeat profiling (Genomics Core Facility at “Alberto Sols” Biomedical Research Institute) and routinely tested for mycoplasma contamination. HL-60 differentiation to macrophages was achieved by adding 1nM of phorbol 12-myristate 13-acetate (TPA) for 24 hours.

### Generation of OSM overexpressing and OSMR knockdown cells

MDA-MB-231 cells were stably transfected with 2 μg of pUNO1-hOSM expression construct (InvivoGen), using TurboFect™ followed by Blasticidin (Sigma-Aldrich) selection at 10 μg/ml. Control transfections were performed simultaneously using 2 μg of empty vector. For OSMR knockdown in CAF-173 cells, pLKO-puro-shOSMR lentiviral vectors were purchased from Sigma-Aldrich (NM_003999.1-1342S21C1). Lentiviral infections were performed as previously described^42^.

### Collagen gel contraction assays

To assess the collagen remodelling capacity^43^, fibroblasts were treated for 4 days with recombinant human OSM (R&D Systems) at 10 ng/μl or vehicle (PBS) before being embedded in collagen (2mg/ml, Corning). After polymerization, collagen gels were detached, and they were treated with OSM (10ng/ml) or vehicle. Pictures were taken 48 hours later, and the area of collagen disks was analysed using Fiji-Image J software.

### 3D fibroblast cell cultures

Fibroblast spheres were formed seeding 8000 cells/well in 96-well ultra-low attachment Corning plates (Costar). Cells were treated with 30 ng/μl OSM or PBS for 3 (for transcriptomic microarray analysis) or 4 days (for RT-qPCR, Western Blot analysis and quantification of spheres area). Pictures were taken using EVOS FL Cell Imaging System (ThermoFisher) and area of spheres was analysed using Fiji-Image J software. Spheres were collected for RNA and protein analysis.

### Mouse studies

All procedures involving animals were performed with the approval of the Biodonostia Animal Experimentation Committee and Gipuzkoa Regional Government, according to European official regulations. Generation of the congenic strain MMTV-PyMT:OSMR−/− was accomplished by mating MMTV-PyMT mice (FVB/N-Tg(MMTV-PyVT)634Mul/J, The Jackson Laboratory) with OSMR^−/−^ mice (OSMR KO, B6.129S-Osmr<tm1Mtan>, Riken BRC)^44,45^. To transfer the OSMR KO line (with a C57BL/6J background) to the genetic background of the tumour-prone animals (FVB/NJ), the OSMR KO mice were previously backcrossed with FVB/NJ mice for 9 generations. Animals used for experiments were female littermates. Tumours were measured once a week using a calliper and volume was calculated as (4π/3) × (width/2)^2^ × (length/2). Animals were culled at 14 weeks, once tumours in the control group reached the maximum allowed size. Tumour burden was calculated by adding the volume or the weight of all the tumours from the same animal. For whole-mount analysis of preneoplastic lesions, abdominal mammary glands from 9 week-old MMTV-PyMT:OSMR−/− and control female mice were spread out on a glass slide, fixed overnight in Carnoy’s solution, stained with Carmine Alum and cleared in ethanol and xylene. Pictures were taken with a Nikon D5000 at 60mm focal length. For the generation of syngeneic orthotopic tumours, 300.000 viable TS1 cells in growth factor reduced (GFR) matrigel (1:1 ratio, Corning), were injected into the fourth right mammary fat pad of anesthetized (with 4% isoflurane) 6-8 week-old OSMR KO or control mice. For the orthotopic co-injections of MDA-MB-231 breast cancer cells and CAF-173 CAFs, cells were injected into the fourth right mammary fat pad of anesthetized (with 4% isoflurane) 6 week-old Athymic Nude-Foxn1nu (Charles River). In OSM activation experiments, CAF-173 were treated with 10 ng/μl OSM for 4 days, prior to co-injection with MDA-MB-231 (500.000 cells each cell line) in GFR matrigel (Corning, 1:1 ratio). For OSMR knockdown experiments, 100.000 MDA-MB-231 hOSM cells and 500.000 CAF-173 shOSMR CAFs were co-injected in GFR matrigel. Animals were monitored 3 times a week and tumour growth measured using a calliper. Animals were culled once tumours reached the maximum allowed size. In all mouse experiments, after animal culling lungs were visually inspected for macroscopic metastases, and mammary glands and lungs were fixed in neutral buffered formalin solution (Sigma-Aldrich). Microscopic metastases were determined by H&E staining of formalin-fixed paraffin-embedded sections. Tumours were divided in portions for 1) preparation of tissue sections for H&E and IHC staining (fixed in formalin) and 2) protein and RNA extraction (snap frozen).

### Flow cytometry sorting

TS1 cells were injected orthotopically in FVB mice as described above, and 15 days after injection, freshly obtained TS1 tumours were dissociated into single cell suspension and stained with the antibodies described in Supplementary Table 1. Flow sorting was performed with a BD FACSAria II cell sorter. Gating strategy for experiments is shown in Supplementary Fig. 1. A pool of 4 tumours from 4 animals were used for each sorting experiment. MMTV-PyMT tumours were sorted by Ferrari *et al*. (2019)^46^ and RNA from FACS sorted tumours was kindly provided by Dr Fernando Calvo. Briefly, tumour populations were separated into fibroblasts (PDGFRA+), cancer (EPCAM+), immune (CD45+), endothelial cells (CD31+), and the remaining population that was negative for all markers.

### Western blotting

Cells and tumours were lysed in RIPA buffer supplemented with protease and phosphatase inhibitors (cOmplete™ ULTRA Tablets, Mini, EASYpack Protease Inhibitor Cocktail, and PhosSTOP, both from Roche). Total lysates were resolved by SDS/PAGE and transferred to nitrocellulose membranes. After blocking with 5% (wt/vol) nonfat dry milk in TBS-Tween, membranes were incubated with the corresponding antibodies (Supplementary Table 1) overnight at 4 °C. Secondary antibodies were chosen according to the species of origin of the primary antibodies and detected by an enhanced chemiluminescence system (Bio-Rad). Densitometric analysis of the relative expression of the protein of interest versus the corresponding control (β-actin or total STAT3) was performed with Fiji-Image J software. Uncropped images used to display blots in the main figures can be found in Supplementary Fig. 2.

### DNA/RNA extraction, RT-qPCR and transcriptomic analysis

Lung genomic DNA was extracted from frozen lungs using the QIAmp DNA mini kit (Qiagen) for qPCR analysis. RNA was obtained from snap frozen animal tissue or cell pellets and extracted using TRIzol reagent (Invitrogen) or Recover all Total Nucleic Acid Isolation kit (Invitrogen), for qRT-PCR and microarray analysis, respectively. cDNA was obtained with the Maxima first strand cDNA synthesis kit (Thermo Scientific) with DNAse treatment incorporated. qPCR was performed using Power SYBR Green PCR master mix (Applied Biosystems), oligonucleotides sequences are described in Supplementary Table 2 and were purchased from Condalab. Expression levels of genes were determined using the ΔΔCt method^47^ and normalized against 3 housekeeping genes optimized for each reaction^48^. Human Alu sequences^17^ were normalized against 18s housekeeping gene capable of recognizing both human and mouse DNA. Microarray analysis was performed using Human Clariom S assay (ThermoFisher). RNA quality was evaluated using the 2100 Bioanalyzer (Agilent) and microarray chips were processed on the Affymetrix GeneChip Fluidics Station 450 and Scanner 3000 7G (Affymetrix) according to standard protocols. Data were analysed using the Transcriptome Analysis Console 4.0 (TAC). Genes with FDR<0.1 and fold change >|2| were considered significantly modulated. GSEA was performed as previously described^49^. FDR < 0.25 or 0.05 was regarded as statistically significant, depending on the type of permutations performed. We compiled the GSEA signatures used in Fig. 3 and 4 and Extended Data Fig. 3 and 4 from the Molecular Signatures Database (MsigDB) by the Broad Institute or they were manually curated from the literature. The gene list for each signature is publicly available at: http://software.broadinstitute.org/gsea/msigdb/search.jsp, Pein *et al.* (2020)^50^ or in Supplementary Tables 3 and 4.

### Histopathology and Immunohistochemistry (IHC) analysis

Histological analysis of murine tumours and lung metastasis was performed in formalin-fixed paraffin-embedded haematoxylin-eosin stained sections. Immunohistochemical staining was performed in formalin-fixed paraffin-embedded sections using the streptavidin–biotin–peroxidase complex method. Antigen retrieval was performed using boiling 10 mM citrate buffer, pH 6.0, for 15 min. Endogenous peroxidase activity was inactivated by incubation with 3% hydrogen peroxide in methanol (15 min, at room temperature). Tissue sections were incubated in a humidified chamber (overnight, 4 °C) using the antibodies described in Supplementary Table 1 diluted in Tris-buffered saline (TBS). For negative controls, primary antibodies were replaced by non-immune serum. After three rinses in TBS (5 min each), samples were incubated with the corresponding secondary antibody (Supplementary Table 1). After 30 min incubation, tissue sections were washed in TBS (5 min, 3 times) and immediately incubated for 30 min with streptavidin–peroxidase complex diluted 1:400 in TBS (Invitrogen). The chromogen was 3-30-diaminobenzidine (Vector Laboratories). Nuclei were counterstained with Harris haematoxylin for 1 min. Pictures were obtained using the Nikon Eclipse 80i microscope with the Nikon DS-5M camera incorporated. The number of positive cells and total cells per area was counted manually in 5-10 different areas of samples from 5 mice per experimental group, using Fiji-Image J software.

### Cytokine and chemokine analysis

Cytokine and chemokine levels were analysed in conditioned media from CAF-173 treated with OSM (30 ng/mL) or vehicle for 72 hours, and from MDA-MB-231-hOSM and corresponding control cells (n = 4 independent experiments). A panel of 31 human chemokines was analysed by Human Chemokine Array Kit (Proteome Profiler Array, R&D Systems), and VEGF levels were quantified by Human VEGF Quantikine ELISA Kit (R&D Systems) following the manufacturer’s instructions. Mouse VEGF, CXCL1 and CXCL16 levels on plasma from 14-week-old MMTV-PyMT: OSMR KO, HET (heterozygous) and WT mice were analysed by mouse Premixed Multi-Analyte Kit (Magnetic Luminex Assay, R&D Systems) following the manufacturer’s instructions. Detection was carried out with the MAGPIX^®^ detector and data analysis was performed using the xPOTENT^®^ software, both from R&D Systems.

### Statistical analyses

Statistical analyses were performed using GraphPad Prism or SPSS softwares. For Gaussians distributions, the student’s t-test (paired or unpaired) was used to compare differences between two groups. Welch’s correction was applied when variances were significantly different. One-way ANOVA was used to determine differences between more than two independent groups. For non-Gaussian distributions, Mann–Whitney’s test was performed. Chi-square test was used to determine differences between expected frequencies. For Kaplan-Meier analysis the Log-rank (Mantel-Cox) test was used. P values inferior to 0.05 were considered statistically significant. Unless otherwise stated, results are expressed as mean values +/− standard errors (SEM).

### Data availability

RNA-seq raw data were obtained from Valdes-Mora *et al.* (2020)^38^ https://www.biorxiv.org/content/10.1101/624890v3. The datasets generated during the current study will be available in the GEO repository upon publication. A confidential reviewer link can be facilitated upon request. Source data on uncropped Western blots are provided in Supplementary Figure 1. The gene list for the fibroblast activation markers signature used in Fig. 3 was derived from Sahai *et al*. (2020)^4^ and is shown in Supplementary Table 3. The gene list for the CAF-173 OSM signature used in Extended Data Fig. 3 includes the 233 genes differentially upregulated in CAF-173 upon OSM treatment and can be found in Supplementary Table 4. Source data for Figs. 1–4 and Extended Data Figs. 1–4 will be provided upon publication and can be facilitated to reviewers upon request. All other data files supporting the findings of this study are available from the corresponding author upon reasonable request.

## Supporting information

Araujo et al Extended Data and Supplementary Information

## Acknowledgements

We are grateful to the members of our laboratories for critical discussion of this work and to the Genomics and Histology Platforms and Animal Facility of the Biodonostia Health Research Institute and Onkologikoa Foundation for technical assistance and advice. We thank Dr Eva González-Suarez (CNIO, Madrid, Spain) and Dr William Muller (McGill University, Montreal, Canada) for providing the MMTV-PyMT mice, and Dr Ander Izeta (IIS Biodonostia, San Sebastian, Spain) and Dr Roger Gomis (IRB, Barcelona, Spain) for providing the HS27 fibroblasts and LM2 cells respectively. This work was funded by Spanish Ministry of Economy and Competitiveness (PI15/00623, PI18/00458 and CP18/00076) and European Regional Development (FEDER) funds, Basque Department of Health (2017111011), Fundación SEOM (Beca SEOM-Font Vella) and Ikerbasque Basque Research Foundation. A.Araujo and A.Abaurrea are funded by Basque Government Doctoral Training Grants and Fundación Gangoiti. JILV is funded by a PFIS Doctoral Training Grant from the Spanish Ministry of Economy and Competitiveness (FI19/00193).

## Author Contributions

A.Araujo, A.Abaurrea, PA and IOQ performed all the cellular and molecular experiments. A.Araujo, A.Abaurrea and MMC performed the animal experiments. JMF performed immunohistochemistry and analysed mouse histopathology. JILV analysed mouse immunostaining. FVM and DGO obtained and analysed the scRNAseq data. A.Araujo, LJ, NF and FC performed FACS-Sorting. AT and SEP collected and analysed patient data and generated the TMAs. RR and MMC analysed the TMA staining. A.Abaurrea, PA, NMM, FC, AC and MMC performed bioinformatic analysis. PFN and PB generated the fibroblast cell lines. NC, IAL and AU contributed with experimental design. CI and CHL contributed with experimental design and helped with supervision of the project. A.Araujo and MMC designed and supervised the study, analysed the data and wrote the manuscript. All authors gave final approval to the submitted and published versions of the manuscript.

## References

1. Hanahan, D. & Coussens, L. M. Accessories to the Crime: Functions of Cells Recruited to the Tumor Microenvironment. Cancer Cell 21, 309–322 (2012).

2. Quail, D. F. & Joyce, J. A. Microenvironmental regulation of tumor progression and metastasis. Nat. Med. 19, 1423–1437 (2013).

3. Lin, E. Y. et al. Progression to Malignancy in the Polyoma Middle T Oncoprotein Mouse Breast Cancer Model Provides a Reliable Model for Human Diseases. Am. J. Pathol. 163, 2113–2126 (2003).

4. Sahai, E. et al. A framework for advancing our understanding of cancer-associated fibroblasts. Nat. Rev. Cancer 20, 174–186 (2020).

5. Shree, T. et al. Macrophages and cathepsin proteases blunt chemotherapeutic response in breast cancer. Genes Dev. 25, 2465–2479 (2011).

6. Curtis, C. et al. The genomic and transcriptomic architecture of 2,000 breast tumours reveals novel subgroups. Nature 486, 346–352 (2012).

7. Wang, Y. et al. Gene-expression profiles to predict distant metastasis of lymph-node-negative primary breast cancer. Lancet 365, 671–679 (2005).

8. Györffy, B. et al. An online survival analysis tool to rapidly assess the effect of 22,277 genes on breast cancer prognosis using microarray data of 1,809 patients. Breast Cancer Res. Treat. 123, 725–731 (2010).

9. West, N. R., Owens, B. M. J. & Hegazy, A. N. The oncostatin M-stromal cell axis in health and disease. Scand. J. Immunol. 88, (2018).

10. Uhlen, M. et al. A pathology atlas of the human cancer transcriptome. Science 357, (2017).

11. Birnie, G. D. The HL60 cell line: A model system for studying human myeloid cell differentiation. Br. J. Cancer 58, 41–45 (1988).

12. Li, T. et al. TIMER: A Web Server for Comprehensive Analysis of Tumor-Infiltrating Immune Cells. Cancer Res. 77, e108–e110 (2017).

13. Aran, D., Hu, Z. & Butte, A. J. xCell: digitally portraying the tissue cellular heterogeneity landscape. Genome Biol. 18, 220 (2017).

14. Koboldt, D. C. et al. Comprehensive molecular portraits of human breast tumours. Nature 490, 61–70 (2012).

15. Fernández-Nogueira, P. et al. Tumor-Associated Fibroblasts Promote HER2-Targeted Therapy Resistance through FGFR2 Activation. Clin. Cancer Res. 26, 1432–1448 (2020).

16. Kucia-Tran, J. A. et al. Anti-oncostatin M antibody inhibits the pro-malignant effects of oncostatin M receptor overexpression in squamous cell carcinoma. J. Pathol. 244, 283–295 (2018).

17. Zijlstra, A. et al. A quantitative analysis of rate-limiting steps in the metastatic cascade using human-specific real-time polymerase chain reaction. Cancer Res. 62, 7083–7092 (2002).

18. Yang, J., Yan, J. & Liu, B. Targeting VEGF/VEGFR to modulate antitumor immunity. Front. Immunol. 9, 1–9 (2018).

19. Jahchan, N. S. et al. Tuning the Tumor Myeloid Microenvironment to Fight Cancer. Front. Immunol. 10, 1611 (2019).

20. Engblom, C., Pfirschke, C. & Pittet, M. J. The role of myeloid cells in cancer therapies. Nat. Rev. Cancer 16, 447–462 (2016).

21. Coffelt, S. B., Wellenstein, M. D. & De Visser, K. E. Neutrophils in cancer: Neutral no more. Nat. Rev. Cancer 16, 431–446 (2016).

22. Daley, J. M., Thomay, A. A., Connolly, M. D., Reichner, J. S. & Albina, J. E. Use of Ly6G-specific monoclonal antibody to deplete neutrophils in mice. J. Leukoc. Biol. 83, 64–70 (2008).

23. Allaoui, R. et al. Cancer-associated fibroblast-secreted CXCL16 attracts monocytes to promote stroma activation in triple-negative breast cancers. Nat. Commun. 7, (2016).

24. Acharyya, S. et al. A CXCL1 paracrine network links cancer chemoresistance and metastasis. Cell 150, 165–178 (2012).

25. Taniguchi, K. & Karin, M. IL-6 and related cytokines as the critical lynchpins between inflammation and cancer. Semin. Immunol. 26, 54–74 (2014).

26. Tower, H., Ruppert, M. & Britt, K. The Immune Microenvironment of Breast Cancer Progression. Cancers (Basel). 11, (2019).

27. Weiner, G. J. Building better monoclonal antibody-based therapeutics. Nat. Rev. Cancer 15, 361–370 (2015).

28. Reid, J. et al. In vivo affinity and target engagement in skin and blood in a first-time-in-human study of an anti-oncostatin M monoclonal antibody. Br. J. Clin. Pharmacol. 84, 2280—2291 (2018).

29. McCarty, K. S. et al. Use of a monoclonal anti-estrogen receptor antibody in the immunohistochemical evaluation of human tumors. Cancer Res. 46, 4244–4249 (1986).

30. Cortazar, A. R. et al. Cancertool: A visualization and representation interface to exploit cancer datasets. Cancer Res. 78, 6320–6328 (2018).

31. Finak, G. et al. Stromal gene expression predicts clinical outcome in breast cancer. Nat. Med. 14, 518–527 (2008).

32. Casey, T. et al. Molecular signatures suggest a major role for stromal cells in development of invasive breast cancer. Breast Cancer Res. Treat. 114, 47—62 (2009).

33. Yeung, T.-L. et al. TGF-β modulates ovarian cancer invasion by upregulating CAF-derived versican in the tumor microenvironment. Cancer Res. 73, 5016–5028 (2013).

34. Nishida, N. et al. Microarray analysis of colorectal cancer stromal tissue reveals upregulation of two oncogenic miRNA clusters. Clin. Cancer Res. 18, 3054–3070 (2012).

35. Calon, A. et al. Dependency of colorectal cancer on a TGF-β-driven program in stromal cells for metastasis initiation. Cancer Cell 22, 571–584 (2012).

36. Mi, H., Muruganujan, A., Ebert, D., Huang, X. & Thomas, P. D. PANTHER version 14: more genomes, a new PANTHER GO-slim and improvements in enrichment analysis tools. Nucleic Acids Res. 47, D419–D426 (2019).

37. Macosko, E. Z. et al. Highly Parallel Genome-wide Expression Profiling of Individual Cells Using Nanoliter Droplets. Cell 161, 1202–1214 (2015).

38. Valdes-Mora, F. et al. Single cell transcriptomics reveals involution mimicry during the specification of the basal breast cancer subtype. bioRxiv 624890 (2020) doi:10.1101/624890.

39. Butler, A., Hoffman, P., Smibert, P., Papalexi, E. & Satija, R. Integrating single-cell transcriptomic data across different conditions, technologies, and species. Nat. Biotechnol. 36, 411–420 (2018).

40. McInnes, L., Healy, J. & Melville, J. UMAP: Uniform Manifold Approximation and Projection for Dimension Reduction. (2018). arXiv:1802.03426v3

41. Guy, C. T., Cardiff, R. D. & Muller, W. J. Induction of mammary tumors by expression of polyomavirus middle T oncogene: a transgenic mouse model for metastatic disease. Mol. Cell. Biol. 12, 954–961 (1992).

42. Etxaniz, U. et al. Neural-competent cells of adult human dermis belong to the Schwann lineage. Stem cell reports 3, 774–788 (2014).

43. Ngo, P., Ramalingam, P., Phillips, J. A. & Furuta, G. T. Collagen gel contraction assay. Methods Mol. Biol. 341, 103–109 (2006).

44. Nakamura, K., Nonaka, H., Saito, H., Tanaka, M. & Miyajima, A. Hepatocyte proliferation and tissue remodeling is impaired after liver injury in oncostatin M receptor knockout mice. Hepatology 39, 635–644 (2004).

45. Tanaka, M. et al. Targeted disruption of oncostatin M receptor results in altered hematopoiesis. Blood 102, 3154–3162 (2003).

46. Ferrari, N. et al. Dickkopf-3 links HSF1 and YAP/TAZ signalling to control aggressive behaviours in cancer-associated fibroblasts. Nat. Commun. 10, (2019).

47. Pfaffl, M. W. A new mathematical model for relative quantification in real-time RT-PCR. Nucleic Acids Res. 29, e45–e45 (2001).

48. Vandesompele, J. et al. Accurate normalization of real-time quantitative RT-PCR data by geometric averaging of multiple internal control genes. Genome Biol. 3, research0034.1 (2002).

49. Subramanian, A. et al. Gene set enrichment analysis: A knowledge-based approach for interpreting genome-wide expression profiles. Proc. Natl. Acad. Sci. 102, 15545 LP – 15550 (2005).

50. Pein, M. et al. Metastasis-initiating cells induce and exploit a fibroblast niche to fuel malignant colonization of the lungs. Nat. Commun. 11, 1494 (2020).

