## Supplementary material for "Stromal Oncostatin M axis promotes breast cancer progression": Araujo et al Extended Data and Supplementary Information

Extended Data Fig. 1

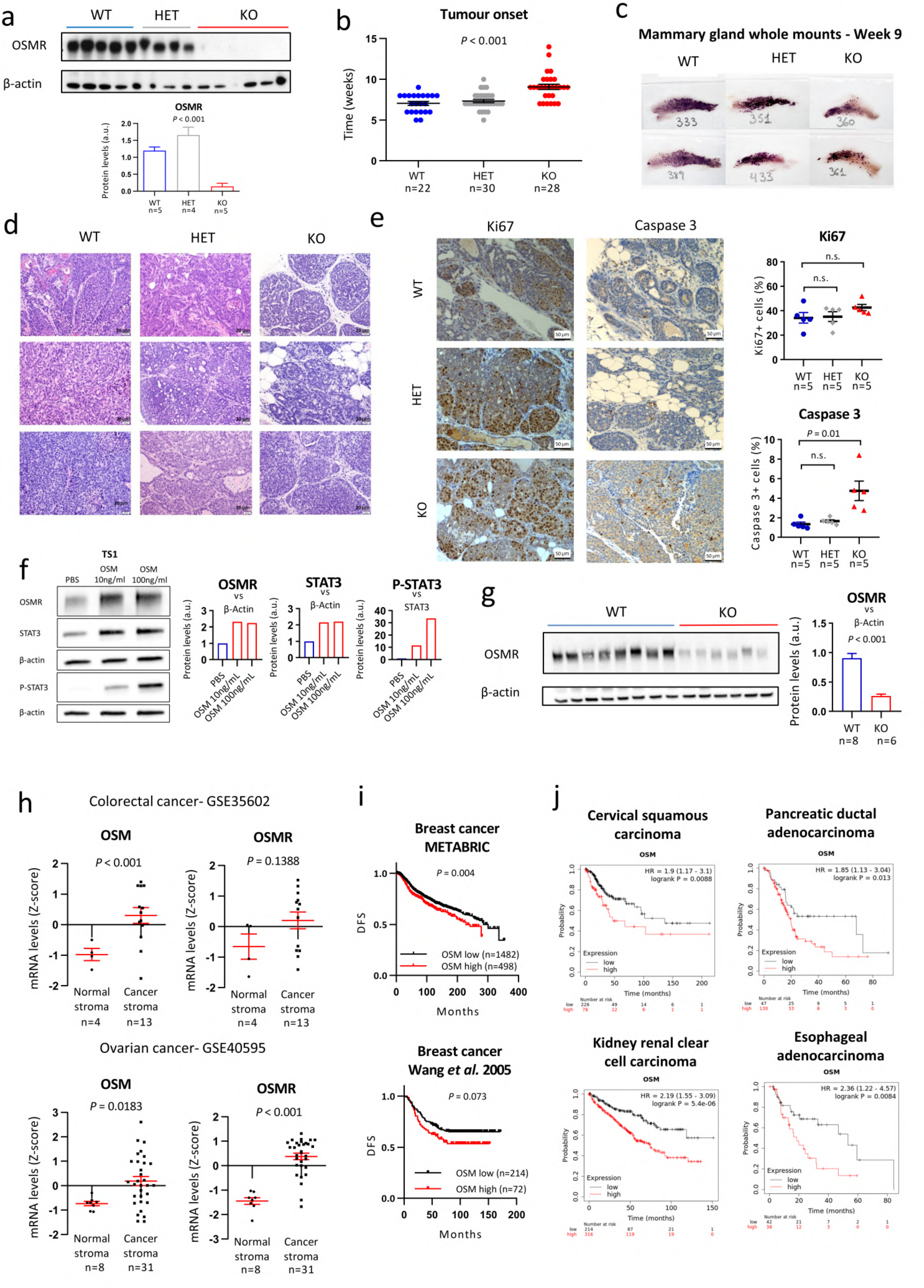

**Extended Data Fig. 1: Effects of OSM:OSMR signalling in cancer progression in murine preclinical models of breast cancer and clinical patients from multiple cancer types. a)** Western blot (upper panel) and densitometric analysis (lower panel) of OSMR protein levels in tumours at culling point from the differential experimental groups of Fig. 1a. **b-d)** Tumour onset **(b)**; representative pictures of whole mount staining of mammary glands at week 9 **(c)** and histopathological analysis of tumours at culling point **(d)** in the different experimental groups of Fig 1a. **e)** Representative pictures (left panels) and quantification (right panels) of Ki67 and active caspase 3 (proliferation and apoptosis markers, respectively) IHC staining in tumours at culling point of the different experimental groups of Fig. 1a. Quantification was performed by manual counting of the percentage of positive tumour cells in a total of 8 pictures per tumour and 5 tumours per group. **f)** Western blot (left panel) and densitometric analysis (right panels) of OSMR, P-STAT3 and STAT3 protein levels in TS1 cells treated with 10 and 100 ng/ml of recombinant OSM for 24h. **g)** Western blot (left panel) and densitometric analysis (right panel) of OSMR protein levels in tumours from animals OSMR WT or KO injected orthotopically with TS1 cells (exp 2, Fig. 1i-l). **h**) OSM and OSMR mRNA expression in normal stroma vs. cancer stroma samples of colorectal and ovarian cancer. Data were downloaded from GEO DataSets (GSE35602 and GSE40595). **i)** Kaplan-Meier curves showing disease-free survival (DFS) for breast cancer patients with high and low OSM expression, included in the METABRIC and Wang datasets. **j)** Kaplan-Meier curves showing overall survival for cancer patients of the indicated tumour type with high and low OSM expression. Data were obtained using KM plotter website. In **i**, **j**) P values were determined using the Mantel-Cox test and high and low OSM levels were determined by automatic cutoff. Unless specified, graphs represent mean ± SEM. P values between the different groups were calculated using one-way ANOVA test **(a,b)** or unpaired two-tailed t test **(e, g, h)**. n.s. Non-significant.

### Extended Data Fig. 2

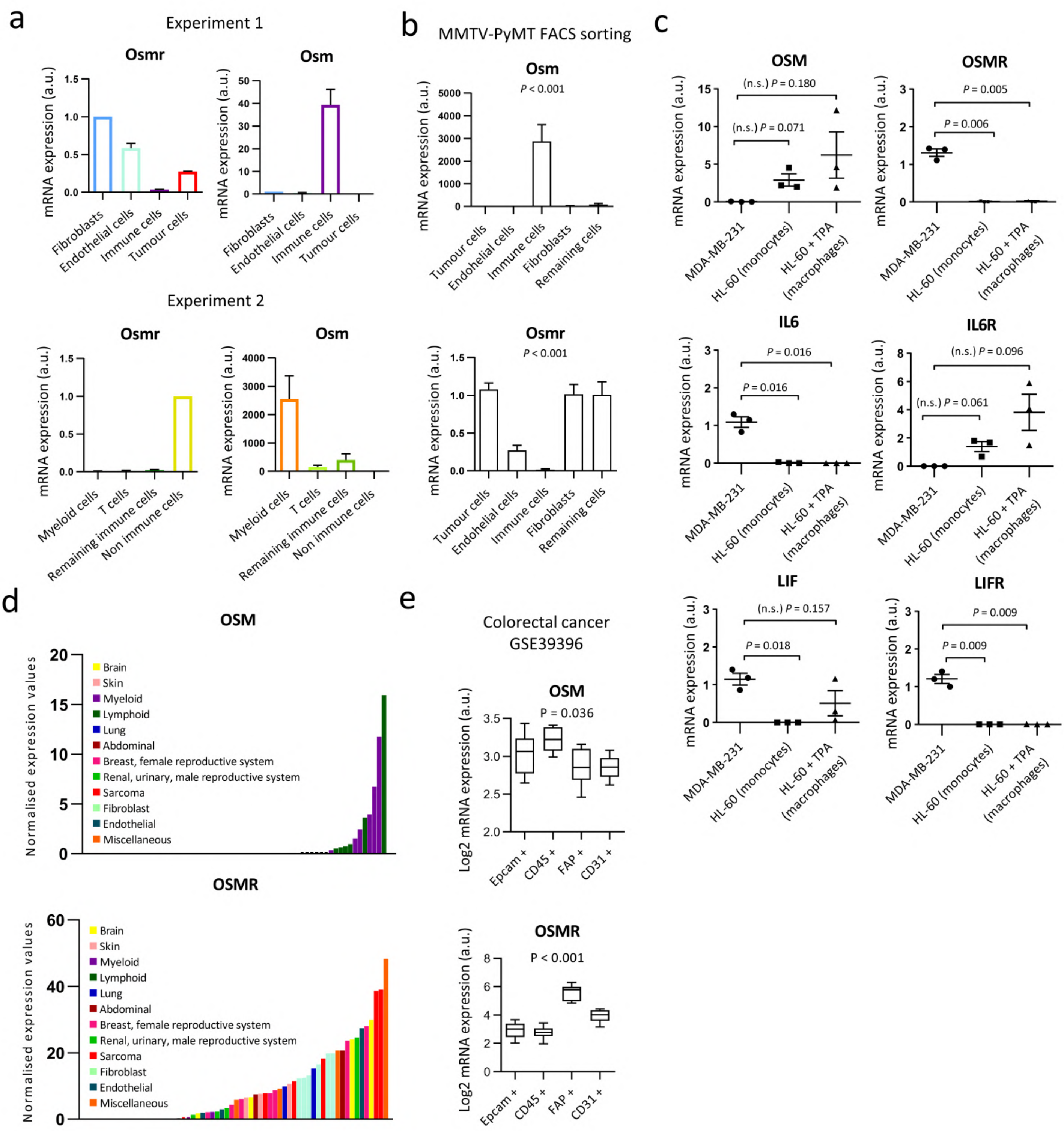

**Extended Data Fig. 2: OSM and OSMR expression in different cell types. a,b)** Osm and Osmr mRNA expression levels analysed by RT-qPCR of FACS sorted populations of TS1 orthotopic tumours **(a)** or MMTV-PYMT FACS sorted tumours described in Ferrari *et al*. (2019) **(b)**. Graphs represent mean ± SEM of 3 technical replicates **(a)** or of 6 different tumours **(b)**. In **a)**, the two different experiments were performed independently, each one in a pool of 4 tumours from individual animals. **c)** mRNA expression levels of the indicated IL-6 family members and associated receptors analysed by RT-qPCR in MDA-MB-231 breast cancer cells and undifferentiated and TPA-differentiated HL-60 cells. Graphs represent mean ± SEM of 3 independent experiments. **d)** OSM and OSMR mRNA relative values in a panel of human immortalized cell lines derived from several body organs. Data were downloaded from Human Protein Atlas. **e**) OSM and OSMR mRNA expression in epithelial cell (Epcam+), immune cell (CD45+), fibroblast (FAP+) and endothelial cell (CD31+) FACS-sorted populations from colorectal cancer samples. Data were downloaded from GSE39396 GEO DataSet. P value was determined using one-way ANOVA test **(b, e)** or the two-tailed unpaired t test **(c)**. n.s. non-significant.

### Extended Data Fig. 3

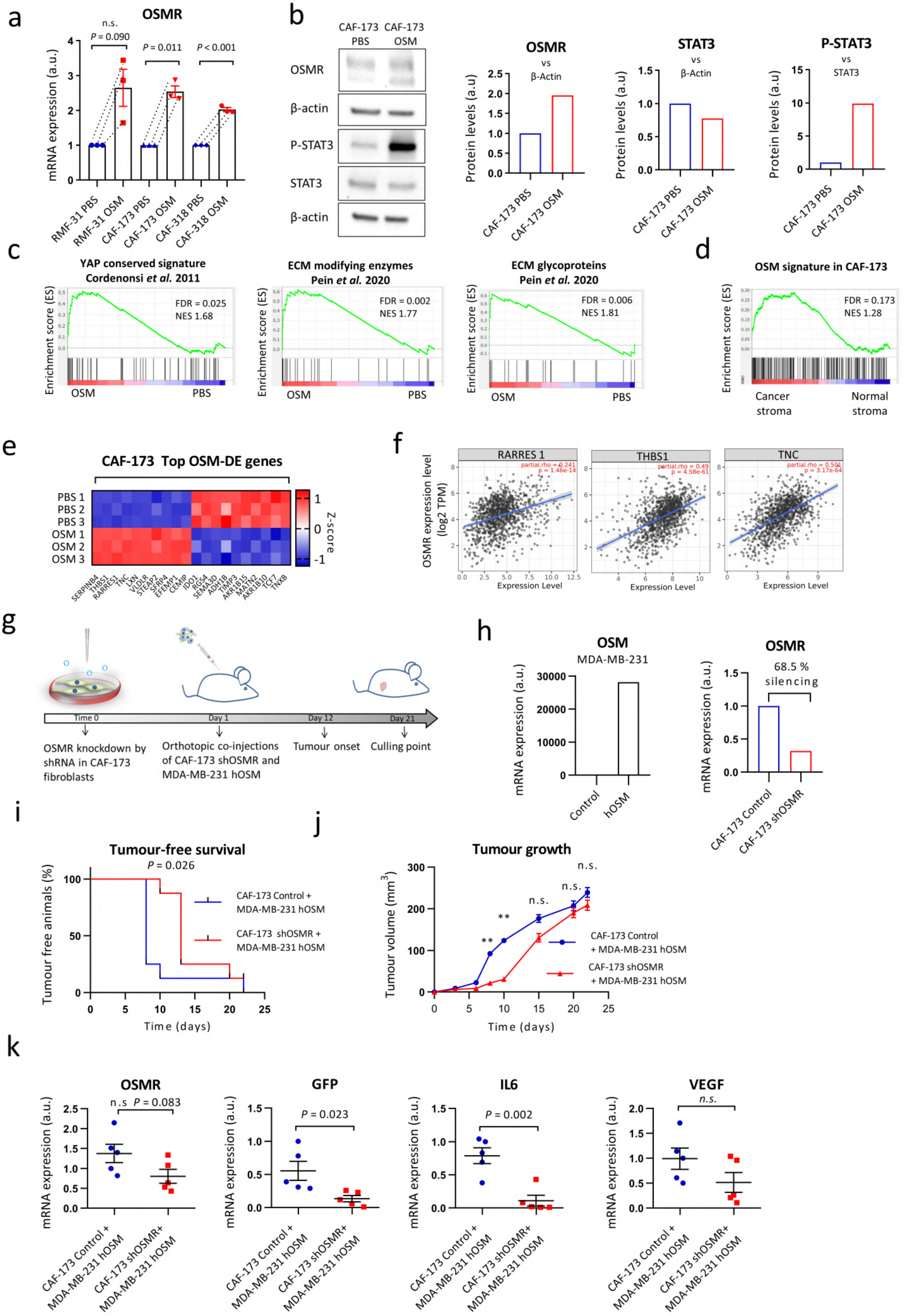

**Extended Data Fig. 3: Effects of OSM in Cancer Associated Fibroblasts (CAFs) *in vitro* and *in vivo*.** **a)** OSMR mRNA expression levels analysed by RT-qPCR in 3D fibroblast spheres treated with OSM for 4 days. n=3 independent experiments. **b)** Western blot (left panel) and densitometric analysis (right panels) of OSMR, P-STAT3 and STAT3 protein levels in CAF-173 treated with OSM. **c)** GSEA showing enrichment of the indicated signatures in microarray data of CAF-173 CAFs treated with OSM. **d)** GSEA showing enrichment of the signature composed of the 233 differentially expressed genes in CAF-173 cells treated with OSM, in microarray data of cancer and normal breast stroma from Finak *et al.* (2008) (GSE9014). **e)** Top 10 up- and down-regulated genes in microarray data of CAF-173 treated with OSM for 4 days. **f)** Correlation of OSMR mRNA levels with RARRES1, THBS1 and TNC expression in breast cancer clinical samples. Data were downloaded from TIMER web platform, Spearman correlation coefficients and P values are shown. **g)** Experimental set-up of the *in vivo* experiment designed to assess the contribution to breast cancer progression of OSMR knockdown in fibroblasts. Control and shOSMR CAF-173 were co-injected with MDA-MB-231-hOSM (500.000 cells each cell line) in matrigel (1:1 ratio) in the mammary gland fat pad of nude mice. n=8 for control CAF-173 + MDA-MB-231 hOSM and n=8 for CAF-173 shOSMR +MDA-MB-231 hOSM. **h)** OSMR and OSM mRNA expression levels analysed by RT-qPCR in MDA-MB-231 and CAF-173 after hOSM and shOSMR plasmid transfection, respectively. **i,j)** Kaplan-Meier curves for tumour-free survival **(i)** and tumour growth **(j)** of orthotopic tumours described in **g)**. **k)** OSMR, GFP, IL6 and VEGF mRNA expression levels analysed by RT-qPCR in 5 tumours per experimental group, as described in **g)**. Graphs represent mean ± SEM. P value was calculated using paired two-tailed t tests **(a)**, the Mantel-Cox test **(i)** or the unpaired two-tailed t test **(j, k)**. n.s. Non-significant. ** p < 0,01.

### Extended Data Fig. 4

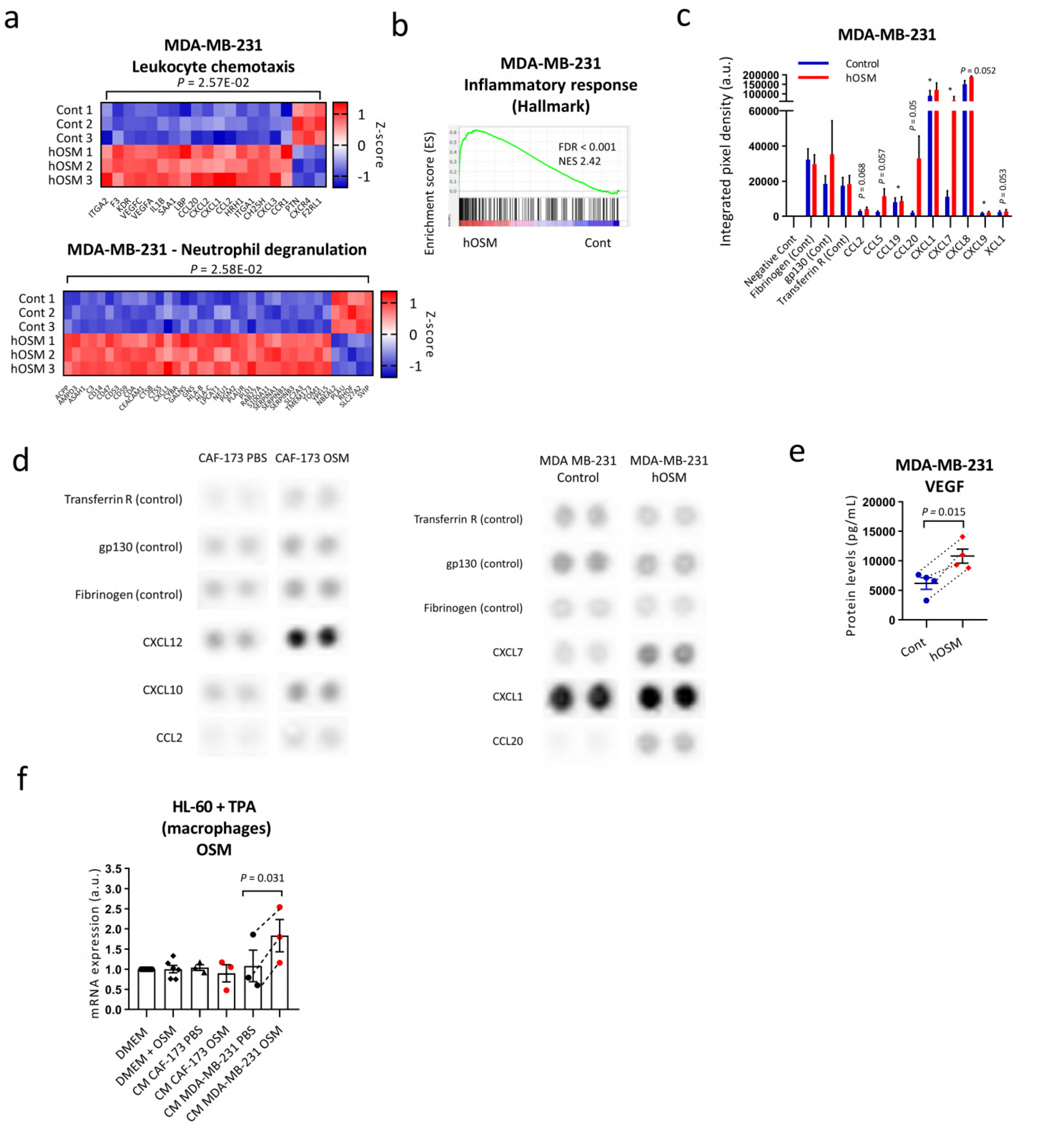


**Extended Data Fig. 4: OSM induces expression of chemokines in cancer cells. a)** Heatmap showing normalized mRNA expression of genes induced by OSM in MDA-MB-231 cancer cells and included in the indicated GO pathways**. b)** GSEA showing enrichment of inflammatory hallmark signature in microarray expression data of MDA-MB-231 hOSM **c, e)** Chemokine array analysis **(c)** and VEGF levels **(e)** of conditioned media from MDA-MB-231-h-OSM and control cells, 72h after seeding. Graphs represent mean ± SEM. n=3 **(c)** or n=4 **(e)** independent experiments. P values were determined using paired two-tailed t tests. **d)** Representative images of chemokine array membranes of conditioned media experiments shown in **c** and Fig. 4c. **f)** OSM mRNA levels in HL-60 + TPA (macrophages) treated with conditioned media from MDA-MB-231 and CAF-173 pre-treated with OSM 10 ng/mL for 72h. Graph represents mean ± SEM of 3 independent experiments. In **c,e,f)**, P values were calculated using the paired two-tailed t test. * p < 0,05.

### Extended Data Fig. 5

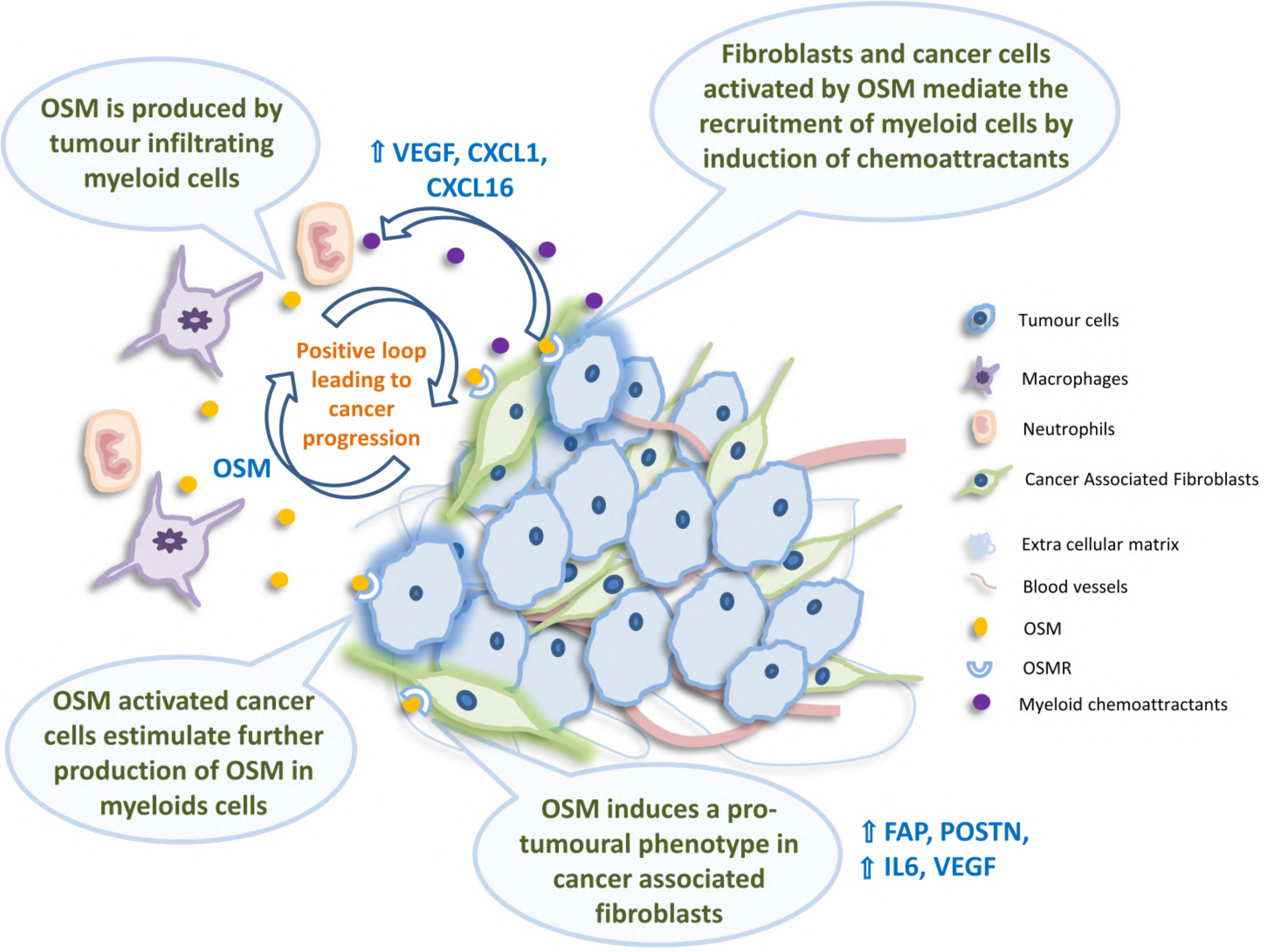


**Extended Data Fig. 5: Proposed working model for the effects of OSM signalling in breast cancer.** Myeloid cells express OSM which activates the OSMR pathway in cancer associated fibroblasts (CAFs). OSM signalling in CAFs induces classical fibroblast activation markers such as FAP, POSTN, IL-6 and VEGF and reprograms CAFs to a pro-malignant phenotype by increasing their contractility and proliferation. At the same time, OSM induces myeloid chemoattractants in CAFs and cancer cells such as VEGF, CXCL1 and CXCL16. This chemokine secretion results in myeloid tumour infiltration, that is further stimulated by cancer cells to induce OSM expression, creating a positive feedback loop that ensures constant secretion of OSM and sustained tumour progression.

Supplementary Figure 1: Experimental design and gating strategy for FACS sorting experiments of Extended Fig. 2a.

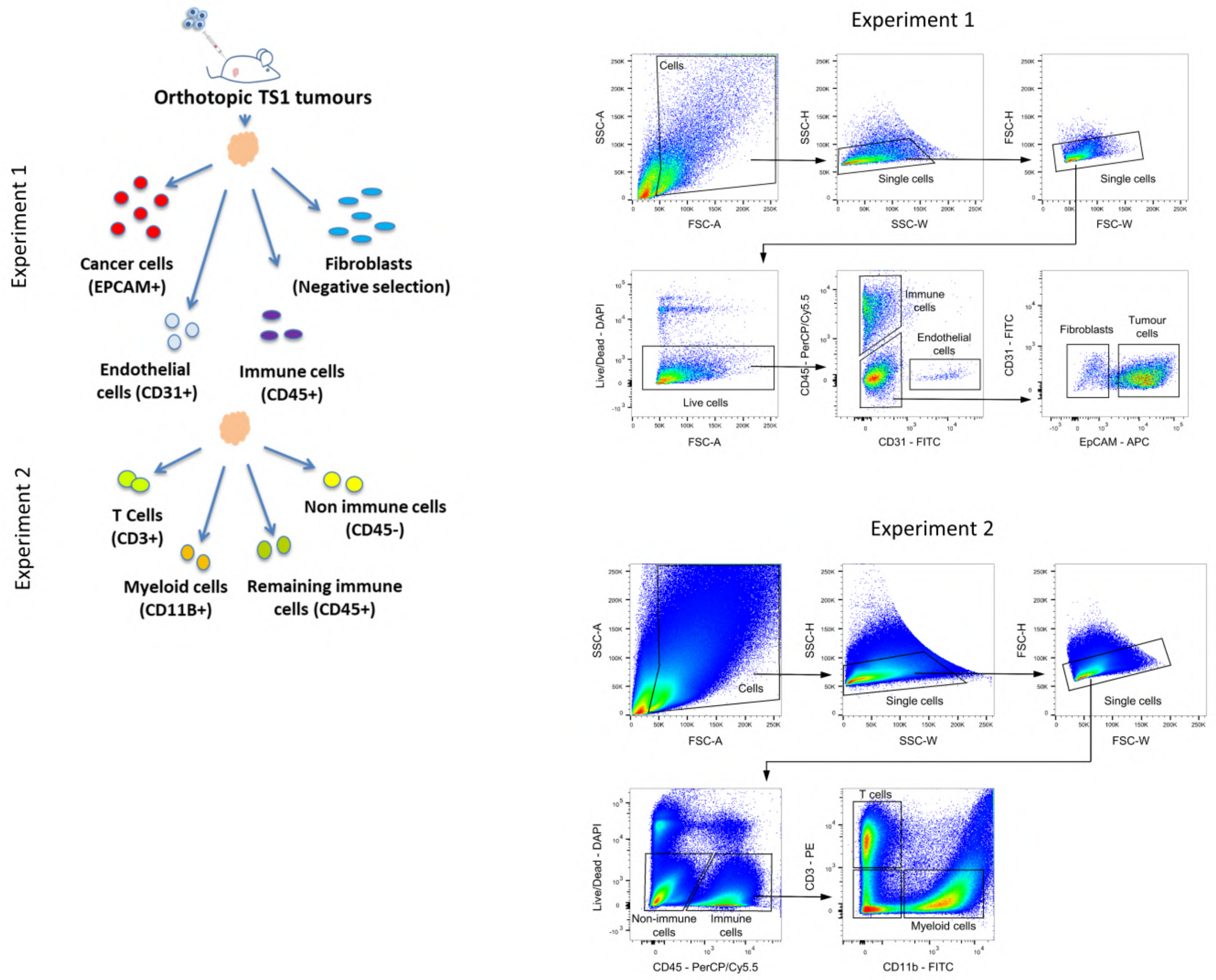

### Supplementary Figure 2: Uncropped Western Blot images for the indicated figures.

Fig. 1f

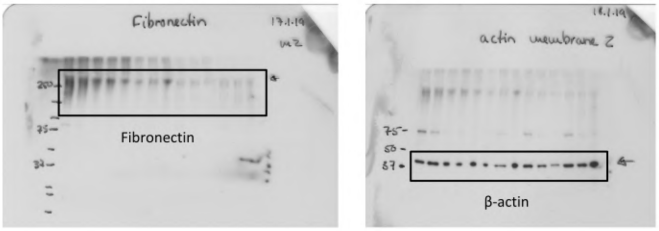


Extended Data Fig. 1a

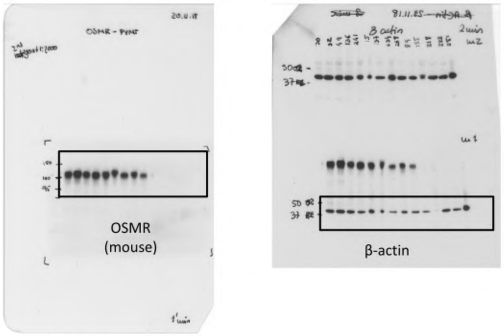


Extended Data Fig. 1f

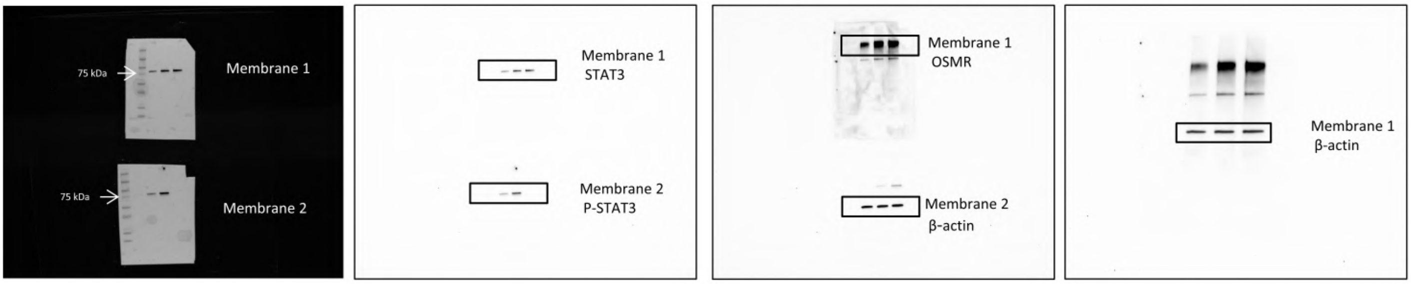


Extended Data Fig. 1g

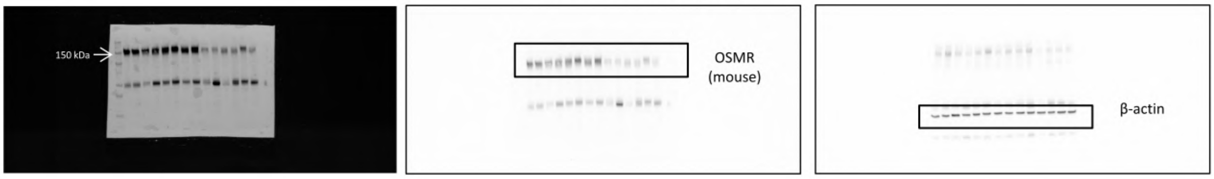


Extended Data Fig. 3b

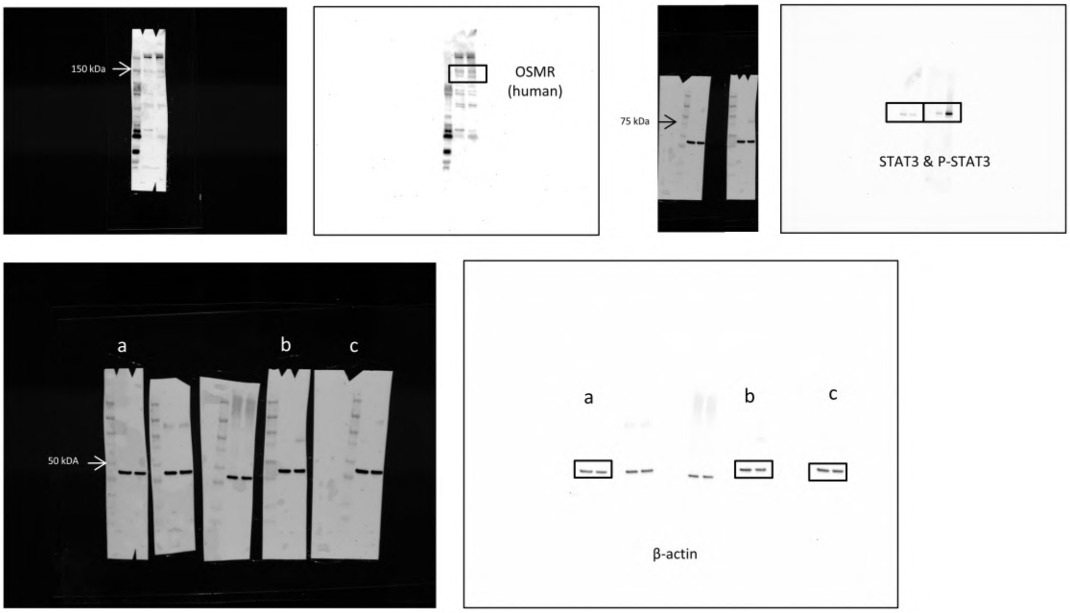

**Supplementary Table 1: Antibodies used in this study.**

| Application | Antibody | Company | Reference | Dilution |
|---|---|---|---|---|
| Western blot | beta-actin (mouse and human) | Sigma | A5441 | 1/2000 |
| Western blot | FN (mouse and human) | Abcam | AB2413 | 1/1000 |
| Western blot | OSMR (mouse) | R&D Systems | AF662 | 1/500 |
| Western blot | OSMR (human) | Santa Cruz | 30010 | 1/500 |
| Western blot | P-STAT3 (mouse and human) | Cell Signaling | 9145 | 1/2000 |
| Western blot | STAT3 (mouse and human) | Cell Signaling | 9139 | 1/1000 |
| FACS | CD11b-FITC | BD Biosciences | 561688 | 1/50 |
| FACS | CD3-PE | Thermo Fisher | 12-0031-82 | 1/40 |
| FACS | CD31-FITC | MACS | 130-097-418 | 1/10 |
| FACS | CD45-PerCP/Cy5.5 | BioLegend | 103131 | 1/80 |
| FACS | EPCAM-APC | Biolegend | 118214 | 1/80 |
| Immunohistochemistry | Caspase 3 | R&D Systems | AF835 | 1/1000 |
| Immunohistochemistry | F4/80 | Bio Rad | MCA497GA | 1/10 |
| Immunohistochemistry | Ki67 | Novocastra | ACK02 | 1/800 |
| Immunohistochemistry | LY6G | BD Biosciences | 551459 | 1/70 |
| Immunohistochemistry | Secondary antibodies | Vector Laboratories Burlingame CA | | 1/400 |

**Supplementary Table 2: qPCR primers used in this study.**

| Gene | Species | Forward primer | Reverse primer |
|---|---|---|---|
| ALU | Human | ACGCCTGTAATCCCAGCACTT | TCGCCCAGGCTGGAGTGCA |
| FAP | Human | CAAAGGCTGGAGCTAAGAATCC | ACTGCAAACATACTCGTTCATCA |
| GP130 | Human | AGGACCAAAGATGCCTCAAC | GAATGAAGATCGGGTGGATG |
| HMBS | Human | GGCAATGCGGCTGCAA | GGGTACCCACGCGAATCAC |
| IL6 | Human | CCAGGAGCCCAGCTATGAAC | CCCAGGGAGAAGGCAACTG |
| IL6R | Human | CCCCTCAGCAATGTTGTTTGT | CTCCGGGACTGCTAACTGG |
| LIF | Human | CCAACGTGACGGACTTCCC | TACACGACTATGCGGTACAGC |
| LIFR | Human | TGGAACGACAGGGGTTCAGT | GAGTTGTGTTGTGGGTCACTAA |
| OSM | Human | CTCGAAAGAGTACCGCGTG | TCAGTTTAGGAACATCCAGGC |
| OSMR | Human | AATGTCAGTGAAGGCATGAAAGG | GAAGGTTGTTTAGACCACCCC |
| POSTN | Human | CTCATAGTCGTATCAGGGGTCG | ACACAGTCGTTTTCTGTCCAC |
| VEGF | Human | AGGGCAGAATCATCACGAAGT | AGGGTCTCGATTGGATGGCA |
| 18S | Human and mouse | CGCGGTTCTATTTTGTTGGT | CGGTCCAAGAATTTCACCTC |

| | | | |
|---|---|---|---|
| HPRT | Human and mouse | TGACACTGGCAAAACAATGCA | GGTCCTTTTCACCAGCAAGCT |
| Actb | Mouse | GCTACAGCTTCACCACCACA | TCTCCAGGGAGGAAGAGGAT |
| Osm | Mouse | ATGCAGACACGGCTTCTAAGA | TTGGAGCAGCCACGATTGG |
| Osmr | Mouse | CATCCCGAAGCGAAGTCTTGG | GGCTGGGACAGTCCATTCTAAA |
| GFP | | CTAGGCCACAGAATTGAAAGATCT | GTAGGTGGAAATTCTAGCATCATCC |

**Supplementary Table 3: Gene list for the fibroblast activation markers signature, manually curated from Sahai et al. 2020, used in Figure 3d.**

| **Fibroblast/ fibroblast subtype/ activation markers** | **Molecules produced by activated fibroblasts/ CAFs** |
|---|---|
| PDGFRa | TGFb |
| Vimentin | VEGFA |
| aSMA | CCN1 (CYR61) |
| FAP | CCN2 (CTGF) |
| Col1a2 | Tenascin |
| Col5a1 | Periostin |
| Downregulation of CD36 | LIF |
| GPR77 | GAS6 |
| CD10 | FGF5 |
| FSP1 (S100A4) | GDF15 |
| | HGF |
| | IL6 |
| | CXCL9 |
| | CXCL12 |
| | FAK |

**Supplementary Table 4: Gene list for the OSM-induced signature in CAF-173 used in Extended Data Figure 3d.**

| | | | | |
|---|---|---|---|---|
| SERPINB4 | PPAP2B | CRISPLD2 | SUSD6 | POSTN |
| THBS1 | STEAP1B | CXCL12 | PFKFB4 | EFCAB13 |
| LXN | CHI3L1 | SERPINB2 | MOSPD1 | B4GALT5 |
| RARRES1 | OSMR | FGF2 | C10orf10 | ABHD17C |
| TNC | SLC2A5 | RHOBTB3 | CTSL | MME |
| VLDLR | GPC6 | ENG | NDST2 | SERPINH1 |
| SFRP4 | TGFBI | FLNB | NPTX2 | TSHZ3 |
| CEMIP | SULF1 | MARCH3 | PRG4 | RGS16 |
| ITGB3 | STEAP1 | GALNT12 | BICC1 | ZDHHC9 |
| MYB | JAK2 | GPC6 | WARS | HOXA4 |
| F2RL1 | IL13RA1 | PUM3 | PCED1A | LAMA4 |
| EFEMP1 | SLC2A14 | LTBP2 | NAMPT | NPC1 |
| KIF26B | ACVR1B | P4HA1 | DUSP1 | VWA5A |
| BRINP1 | CCDC71L | CPD | RUNX2 | POLR2H |
| SERPINE1 | SPRY1 | NAV1 | POLM | SMG6 |
| STEAP2 | MASP1 | TRIB2 | PREB | HEG1 |
| GJA1 | PDK1 | ADAM19 | PTGS2 | GLT8D2 |
| DHRS3 | NREP | WISP1 | SLC16A3 | PTX3 |
| SEL1L3 | PDP1 | ARRDC4 | CYR61 | C1R |
| MYC | SLC2A3 | PTPN2 | RNF144A | SYTL2 |
| IL1R1 | GYS1 | ZPLD1 | NXF3 | SNAI1 |
| RDH10 | TMTC2 | XYLT1 | ADAMTS4 | CDKN1A |
| SERPINB7 | NRP2 | KCTD10 | ADAM12 | TGFBR1 |
| COL5A1 | PCBP3 | GPAM | CLCA2 | ERO1A |
| PLOD2 | BCL6 | RP11-351M8.2 | PGM3 | DLC1 |

| | | | | |
|---|---|---|---|---|
| FGF7 | ADM | C1QTNF6 | LCE2A | JAK2 |
| SERPINB3 | IL6 | CTGF | SLC22A23 | DENND1B |
| LRRC15 | MGP | SNX10 | IGF1R | FHL2 |
| WWC1 | SDC1 | BGN | BNIP3 | ALDH1L2 |
| NID2 | FAM20C | FCHSD2 | TMEM45A | CFAP54 |
| ITGB8 | MTHFD2 | DAB2 | JUNB | DNMT3B |
| NNMT | OSR1 | TXNIP | FNIP2 | SLC16A7 |
| HMOX1 | RAI14 | GABRE | MICAL2 | OPN3 |
| VEGFA | RCAN2 | RCL1 | SMYD3 | SRPX |
| C2orf83 | AK4 | FAM60A | HAS2 | DMGDH |
| SOCS3 | PFKFB3 | IL4R | HS3ST3A1 | BHLHE40 |
| PRDM1 | SBNO2 | ELL2 | TNS3 | GK |
| ARHGAP20 | NABP1 | F3 | CCL2 | CAMK2N1 |
| DDIT4 | PLPP3 | LOX | GALNT2 | RASD1 |
| COLEC12 | GLIS3 | COL5A2 | SMIM3 | NTNG1 |
| TMOD1 | TUBB3 | AGTRAP | SOS1 | ADAMTS3 |
| HS3ST3B1 | PLPP4 | FAM20A | PAPPA | HLA-DQB1 |
| ARL4C | PVR | AGFG1 | GFPT2 | KLF9 |
| IL33 | PGK1 | PDGFRA | NRXN3 | EPB42 |
| FNDC1 | THSD4 | THBS2 | CTBP2 | STX19 |
| TAGLN | HIF1A | COL8A1 | C1orf158 | |
| TNFRSF10D | NCAM2 | PLAT | TMEM2 | |
